# Noninvasive sleep monitoring in large-scale screening of knock-out mice reveals novel sleep-related genes

**DOI:** 10.1101/517680

**Authors:** Shreyas S. Joshi, Mansi Sethi, Martin Striz, Neil Cole, James M. Denegre, Jennifer Ryan, Michael E. Lhamon, Anuj Agarwal, Steve Murray, Robert E. Braun, David W. Fardo, Vivek Kumar, Kevin D. Donohue, Sridhar Sunderam, Elissa J. Chesler, Karen L. Svenson, Bruce F. O’Hara

**Author notes:** These authors contributed equally (Shreyas S. Joshi, Mansi Sethi). Address for correspondence and proofs: Shreyas S. Joshi, Ph.D. Dept. of Biology University of Kentucky 675 Rose Street 101 Morgan Building Lexington, KY 40506 U.S.A.

## Abstract

Sleep is a critical process that is well-conserved across mammalian species, and perhaps most animals, yet its functions and underlying mechanisms remain poorly understood. Identification of genes and pathways that can influence sleep may shed new light on these functions. Genomic screens enable the detection of previously unsuspected molecular processes that influence sleep. In this study, we report results from a large-scale phenotyping study of sleep-wake parameters for a population of single-gene knockout mice. Sleep-wake parameters were measured using a high throughput, non-invasive piezoelectric system called PiezoSleep. Knockout mice generated on a C57BL6/N (B6N) background were monitored for sleep and wake parameters for five days. By analyzing data from over 6000 mice representing 343 single gene knockout lines, we identified 122 genes influencing traits like sleep duration and bout length that have not been previously implicated in sleep, including those that affect sleep only during a specific circadian phase. PiezoSleep also allows assessment of breath rates during sleep and this was integrated as a supplemental tool in identifying aberrant physiology in these knockout lines. Sex differences were evident in both normal and altered sleep behavior. Through a combination of genetic and phenotypic associations, and known QTLs for sleep, we propose a set of candidate genes playing specific roles in sleep. The high “hit rate” demonstrates that many genes can alter normal sleep behaviors through a variety of mechanisms. Further investigation of these genes may provide insight into the pathways regulating sleep, functional aspects of sleep, or indirect potentially pathological processes that alter normal sleep.

## INTRODUCTION / BACKGROUND AND SIGNIFICANCE

Sleep is a complex behavior common to all birds and mammals, and probably most or all other vertebrates and invertebrates with a nervous system. Regulated by a multitude of neuronal processes and indirectly by gene networks, it is a process vital for an organism’s wellbeing. Sleep has been suggested to have a role in learning, memory consolidation, energy restoration, synaptic optimization and recently it has also been implicated in the clearance of potentially toxic metabolites, including β-Amyloid (1–7).

Genetic manipulations have advanced our knowledge about some aspects of sleep, including influences on the sleep EEG, sleep disorders, brain areas regulating sleep processes, and molecular pathways underlying sleep and its regulation. However, many of the molecular mechanisms and pathways underlying sleep remain unknown, and there are still many unresolved questions regarding the biological need for sleep, functions of sleep, and the genetic and physiological basis of sleep homeostasis (8–10). There have been numerous efforts to address these questions utilizing a variety of animal models including mice. These range from individual labs studying specific knockout mice, to large-scale QTL (quantitative trait loci) and genome-wide approaches including phenotype-driven ENU (N-ethyl-N-nitrosourea) mutant screens involving many labs, and gene-driven knockout mouse phenotyping programs (11). Several gene mutations or gene knockouts in mice have been examined for hypothesized effects on sleep. The discovery of novel sleep mechanisms and pathways can be addressed with a large scale screening approach.

Circadian clock genes such as *Clock* and *Rab3a* in mice, and *Per,* and *Dbt* in flies, which influence both circadian timing and sleep homeostasis, have been identified using ENU/EMS mutagenesis techniques (10,12–15). Discovery of these genes led to identification of many others (*Bmal1/Cyc, Cry1,2, Per1,2,3,* etc.) that also underlie circadian and homeostatic aspects of sleep (16). However, many of these mutations produce only subtle phenotypes, which are difficult to detect, and are often affected by the genetic background of the mouse (17). Approaches using traditional mouse strains, genetic crosses, and quantitative trait locus (QTL) mapping strategies have also identified a modest number of sleep-influencing genes such as *Homer1a*, *Acads* (acyl-coenzyme A dehydrogenase), and *Rarb* (Retinoic acid receptor beta) (18–20).

A major bottleneck in large-scale genetic studies of sleep is the difficulty, expense and time demands of traditional EEG/EMG studies. While knockout studies of select target genes such as neurotransmitter receptors have found modest effects on at least one sleep parameter, relatively few genes have been examined (21). Using a higher throughput, non-invasive approach allows for much larger numbers of mice to be examined (22–26). The PiezoSleep system utilizes a sensitive piezoelectric film covering the mouse cage floor, and is especially well suited to characterization of large-scale resources such as mice from the International Knockout Mouse Consortium (IKMC) (27). IKMC aimed at generating mutant embryonic stem cells (ES) with loss of function for each coding gene in the mouse genome on the B6N background. Continued generation of these single-gene KO lines is now being carried out using CRISPR methodologies. As live mice are bred from these approaches, the single-gene knockouts undergo a core set of broad-based phenotyping screens at the KOMP^2^ centers and as part of the International Mouse Phenotyping Consortium (IMPC) (28–31). A collaborative phenotyping approach like KOMP^2^ has many advantages over individual lab efforts, as a single mouse can be examined for multiple traits, and uniform quality control procedures help to limit confounding. In addition, a collective study of these traits allows for associations to be identified that might reveal previously unknown relationships among phenotypes. This broad-based platform aims to expedite the functional annotation of genes, especially those that are currently most poorly understood (32–34). The Jackson Laboratory KOMP2 Phenotyping Center (JAX-KOMP^2^) uniquely integrated PiezoSleep as part of the pipeline, and the results to date are described in this report.

## METHODS

### Generation of KO mice

IKMC mouse mutants were generated on a C57BL/6N mouse background and have either a single null mutation in which an entire locus is removed, or “knockout-first” alleles which permits generation of conditional alleles by utilization of site-specific recombinase (35,36). For phenotyping at JAX KOMP^2^, C57BL/6N is used as the reference control strain and will be referred to as B6N in this report.

As part of the JAX-KOMP^2^ Phase One phenotyping pipeline, each mouse was characterized in a standardized protocol from age 4-18 weeks. During this period, data for more than 200 parameters were collected using a battery of assays. These assays cover a range of morphological, physiological and behavioral traits including many disease relevant parameters pertaining to neurobehavior, metabolism, immune, cardiovascular, sensory, and musculo-skeletal systems, followed by terminal collection of blood and histopathology (37) (Supplementary Figure 1). Additional tests such as light/dark box and hole-board exploration tests, rotarod, and sleep, were unique to the JAX-KOMP^2^ pipeline. Sleep was evaluated using a PiezoSleep System (*Signal Solutions, LLC, Lexington, KY*), a non-invasive, high throughput sleep-wake monitoring system (details provided in following sections). The primary traits analyzed in this system are total sleep duration averaged across 24 h, 12 h light phase and 12 h dark phase, average sleep bout lengths (across 24 h, 12 h light phase, and 12 h dark phase), and breath rate during sleep.

**Figure 1:**
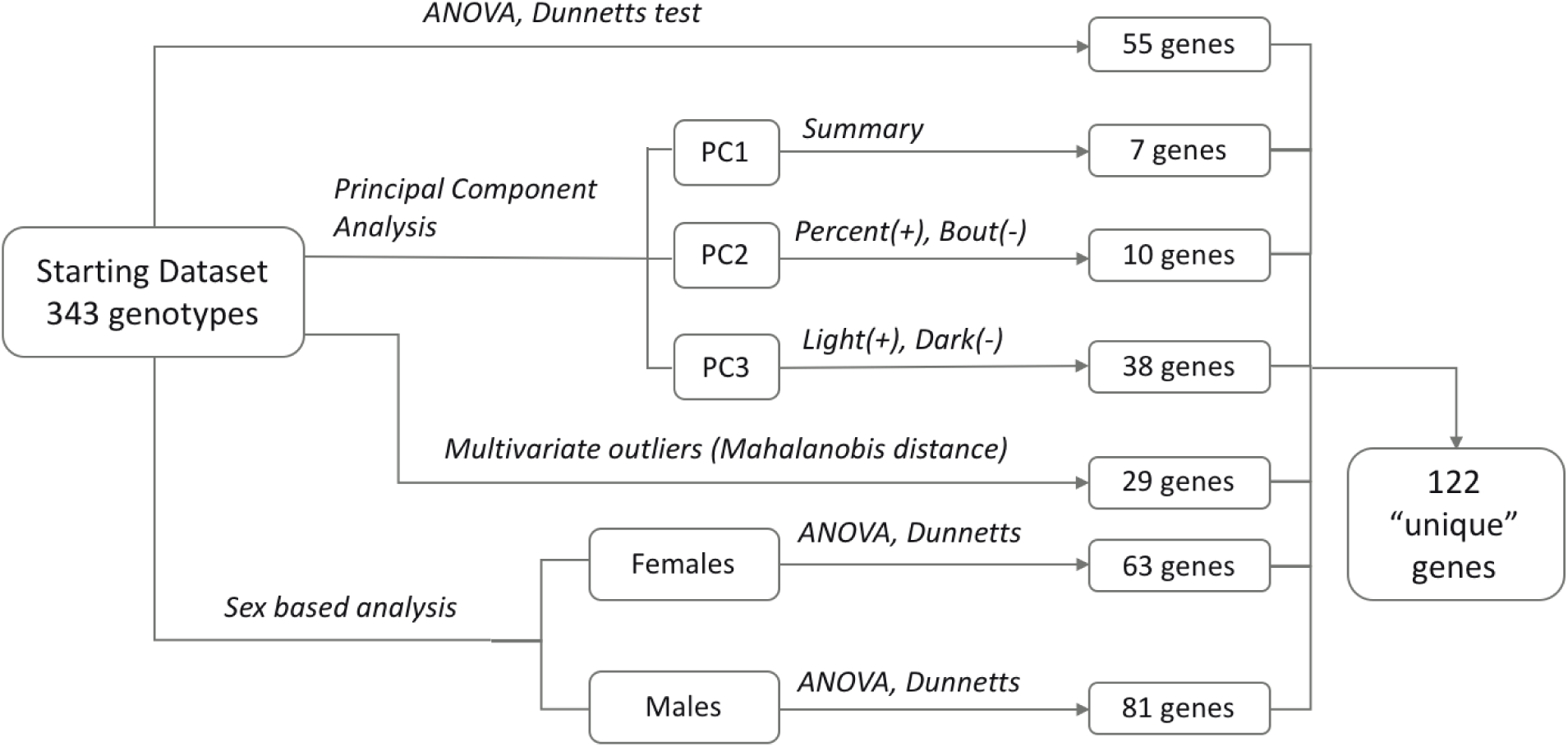
Schematic of data analysis procedure used to derive a set of priority candidate genes.

### Sleep recording with the piezoelectric system

Sleep and wake states were determined using the PiezoSleep System; a quad-cage and piezoelectric sensor system comprised of Plexiglas cages lined with piezoelectric films across the entire cage floor (*Signal Solutions LLC, Lexington, KY*). The piezo system has been validated against simultaneous EEG-based human scoring, and gives a classification accuracy of over 90% for mice (22–24). The piezoelectric films are highly sensitive to pressure changes caused by any movement. During sleep, breathing is the dominant movement, which further allows for an accurate recording and assessment of breath rates. Sleep is characterized by quasi-periodic signals with low variations in amplitude, whereas wakefulness (both active and resting) is characterized by irregular transient and high amplitude pressure variations corresponding to voluntary body movements and weight shifting. Even during quiet rest, the system is capable of detecting subtle head movements associated with olfactory sampling that distinguish this type of wake from true sleep.

To differentiate between sleep and wake states, signal features sensitive to the differences between these states are extracted from 8-second segments, and classified automatically every 2 seconds through overlapping windows. Sleep-wake decisions in the 2-second intervals are binned over specified time periods (e.g., 12 hr, 24 hr) to calculate percent sleep/wake statistics. The system is flexible and allows users to specify their own time intervals (e.g. 5 min, 1 h, 12 h) to compute sleep/wake percent in that duration. In addition, durations of uninterrupted runs of sleep state labels are used to compute mean sleep bout lengths. To eliminate the impact of short and ambiguous arousals on the bout length statistic, a bout length count is initiated when a 30-second interval contains greater than 50% sleep and terminates when a 30-second interval has less than 50% sleep. This 30-second interval or window can be set to shorter time periods, which will then give shorter average bout lengths, but the relative differences between strains typically do not change. In this report, we have used the 30-second window.

### Animal housing and phenotyping

This study utilized control and KO mice produced in the same vivarium. Mice of weaning age were housed at 3-5 mice per pen in pressurized, individually ventilated cages using pine shavings as bedding, with free access to acidified water and food (LabDiets 5K52, LabDiet, Scott Distributing, Hudson, NH). The housing facility was maintained on a 12:12 light/dark cycle starting at 0600 hrs. At 15 weeks of age, mice were removed from their home cages and placed into individual piezo sensor cages. The system used in the JAX-KOMP2 pipeline comprises twenty quad-cage units, allowing for simultaneous assessment of up to 80 mice for a week-long experiment. During the PiezoSleep assay, the light cycle and food and water access were essentially identical to that of standard housing conditions, with normal pine shavings as bedding, which does not significantly alter the piezo signal. In each testing week, 10 age-matched control B6N mice (five females and five males) and 3-19 mice KO mice were tested. Mice were housed in the sleep system for 5 days. For total sleep times and bout lengths across each 24 h period and across each 12h light and dark period, we averaged three consecutive days beginning at light onset of day 2.

### Data Analysis

We began with a dataset containing observations from 8849 mice (4446 females, and 4403 males). In the initial screening step, observations were included from only those mice that were verified to have completed the phenotyping pipeline. One of the outputs of PiezoSleep is a data confidence metric that ranges from 0 through 1 to assess signal quality and potential outliers. Sleep recordings with a data confidence value below the threshold of 0.6 were excluded from the analysis. Control mice were also screened for outliers based on extreme high/low values for their sleep/wake parameters. This was achieved by carrying out multivariate outlier analysis and computing the Mahalanobis distance (md) (38), which is a variance-normalized multidimensional distance of each of the observations from the centroid (vector mean) of all measured variable scores (39). At a default quantile threshold of 0.975, a cutoff value of 4.0 was generated with 114 control mice having md values above the cutoff, and were thus excluded from the analysis. The final dataset used for analysis contained 6350 mice belonging to 343 KO strains and control mice. This final dataset consisted of 1884 Control (918 Females; 894 Males), and 4466 KO (2250 Females; 2216 Males) mice. One-way analysis of variance (ANOVA) was performed for each sleep variable and breath rate, and posteriori multiple comparisons were done using Dunnett’s test to identify genotypes showing significant difference(s) with respect to the control group. Similar analysis was done to identify genotypes with sex-specific effects. 289 of the knockout cohorts containing at least 3 females and 3 males each were included in this analysis. Significance values for simple and interaction effects between genotype and sex were computed through multifactorial ANOVA and effect sizes were estimated. For all of the analyses of the sleep-wake parameters mentioned thus far, an adjusted p-value of less than 0.05 was considered significant.

Principal component analysis (PCA) was performed on six of the sleep variables under consideration: sleep durations and bout lengths averaged across each 24-hour period, and independently for each 12 h light and dark period independently (with the first acclimation day excluded). PCA is a method of multivariate analysis through dimension reduction: i.e., it is used to reduce a large set of variables into principal components (PC) that account for most of the variance of the original variables. The first principal component, if it accounts for most of the variance in the data, can be considered to adequately summarize the data. Analysis of genotype effects on the first PC was performed using ANOVA and the Dunnett’s post-hoc test to identify gene deletions that significantly affect overall sleep. In the final step, mean values for each sleep variable from the piezo system were computed for all genotypes and multivariate Mahalanobis distance outlier analysis was performed for the whole dataset. The genes identified as outliers were considered to have the greatest effect on sleep variables. A final set of all significant genes from all analyses, considered priority candidate genes affecting sleep in mice, was derived from this process (Figure 1).

To investigate relationships between results from our analysis and some of the known circadian and sleep-influencing genes, we conducted an association analysis using GeneMANIA, a tool for analyzing sets of genes (40). The Cytoscape software for network visualization contains a dedicated plugin for GeneMANIA and was used in this analysis. We finished the analysis by identifying other phenotypes associated with sleep. To identify these phenotype associations, data was collected for the final list of sleep-related candidate genes from the IMPC database (mousephenotype.org) through batch query.

GeneWeaver software was used to perform gene set similarity analysis (41). Data analysis was performed in R programming environment in the R Studio (42). Outlier analysis was done using Chemometrics package, and Dunnett’s test was performed using the DescTools package (43,44). The package ggplot2 was used to prepare plots(45).

## RESULTS

### B6N Control mice

We found that female B6N have a significant reduction in total sleep time (*t-test, p<0.001*), as well as reduced sleep in both the light and dark periods (Figure 2A) compared to B6N males, though it is less pronounced during the light phase (*t-test, p < 0.01*) as compared to dark phase (*t-test, p<0.001*). The mean percent sleep across 24 h was 41.83 ± 0.12% for females and 45.11 ± 0.12% for males. During the dark phase, mean sleep duration was 21.53 ± 0.17% in females and 27.21 ± 0.18% in males, and in the light phase, for females sleep duration was 62.13 ± 0.16 % and 63.01 ± 0.15% in males. Similar to sleep duration patterns, females also show shorter bout lengths measured across 24 h (Females: 369.68 ± 2.06 s; Males: 444.78 ± 2.55 s; *t-test, p<0.001*) (Figure 2A), and during both the dark (Females: 198.57 ± 1.44 s; Males: 275.85 ± 2.2 s; *t-test, p<0.001*) and light phases (Females: 539.29 ± 3.03 s; Males: 619.75 ± 3.55 s; *t-test, p<0.001*) (Figure 2B). Overall, B6N female mice have reduced sleep duration and shorter bout length than their male counterparts.

**Figure 2:**
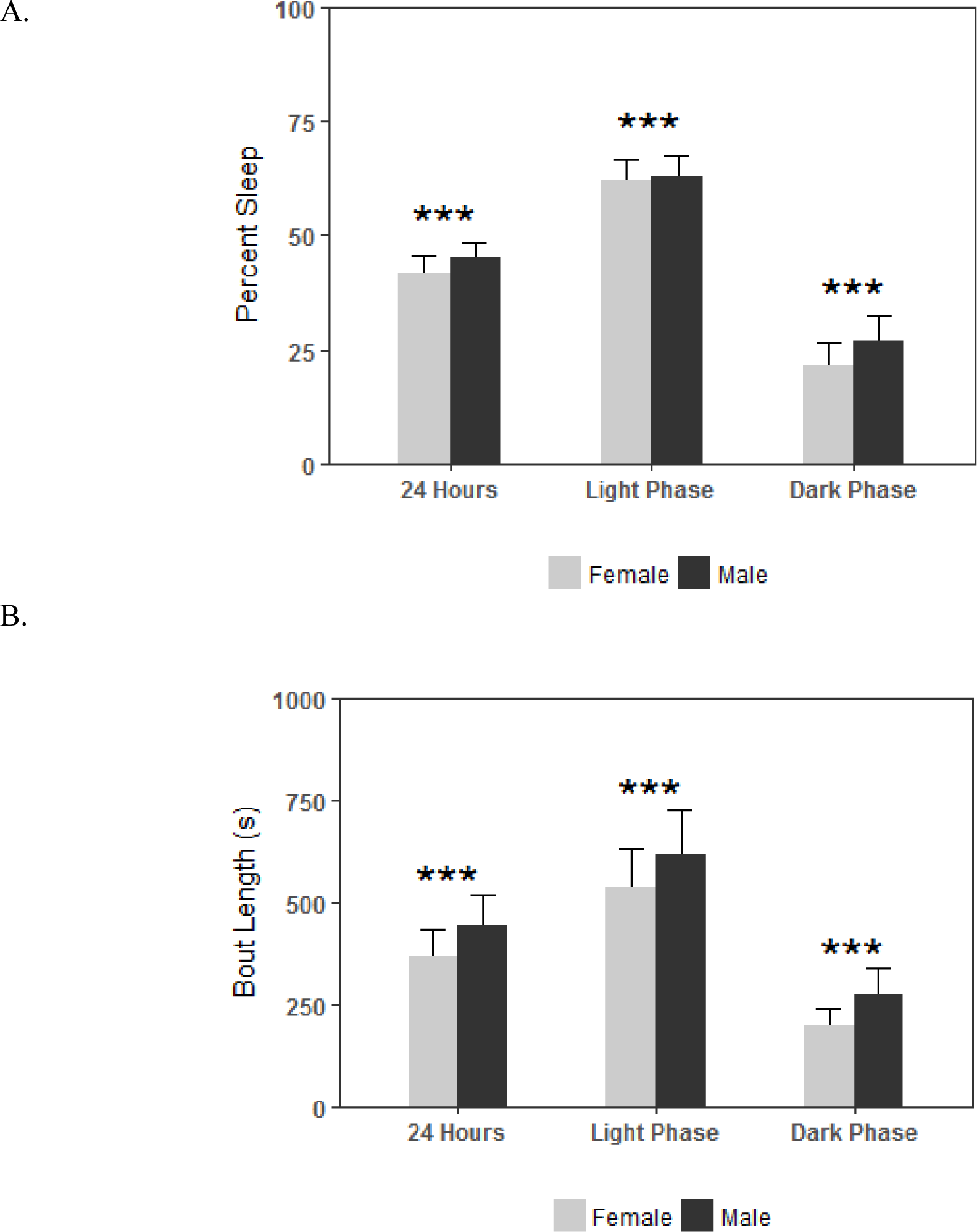
Sleep–wake patterns in control B6N mice. Average percent sleep across three consecutive days (beginning on day 2) and analyzed for (A) 24 h, light phase, and dark phase. Female mice show reduction in sleep duration across 24 h and during the light and dark phase. (B) depicts average bout length in seconds (s) over 24 h, light phase, and dark phase. Females had shorter average bout lengths across all phases. Values represent mean ± SD. ***P < 0.001 from *t-test*.

### Knockout mice

This analysis includes piezo system sleep recordings from 343 KO strains. Of these, 55 (16%) showed significance at *p<0.05* (adjusted for FWER) in Dunnett’s post hoc analysis for one or more of the sleep variables recorded (Supplementary Figure 2). Across the 24-hour period, reduction in sleep percent (total sleep) was observed in *Ppp1r9b, Pitx3, Ap4e1, Cdc20, Vsig4,* and *Bzw2* KO strains, while increased total sleep was seen in *Cbln3, Tpgs2,* and *Ccl26* compared to control mice. During the light phase, *Pitx3, Ppp1r9b, Hsd17b1, Tmem79, Bzw2, Nrcam, Kcnh3,* and *Htr3b* exhibited reduced sleep percentages while longer sleep duration was found in *Eogt, Gjd4, Ccl26, Epgn, Ncald, Neurl2, Npm3, Serpinb5, Htr1f, Ifnl3, Foxo3, Dnaja4, Cpb1, Bex4, Ttll6, Rimklb, Ydjc, Zfp961, Slc46a3,* and *Slc8b1* KO strains. During the dark phase, sleep duration was reduced for *Ppp1r9b, Vsig4, Rimklb,* and *Mylip*, and increased for *Cbln3, Macrod2,* and *Postn*. Additionally, as compared to controls, mean bout length was significantly reduced across 24h in *Pitx3, Hsd17b1, Myh1, Rnf10, Myo3b, Ap4e1,* and *Ppp1r9b*, and increased in *Tmem136* mutant mice. In light phase, bout lengths were significantly shorter in *Pitx3, Hsd17b1, Ptpru, Myh1, Ppp1r9b, Ap4e1,* and *Nrcam* and longer in *Arrb2, Adck2, Slc8b1, Zfp961, Htr1d, Zbtb4, Tmem136, Emp1,* and *Ipp.* During dark phase, significantly shorter mean bout lengths were seen in *Stx16, Rimklb, Rnf10, Myo3b, Bex4, Ppp1r9b, Tmem151b, Zzef1, Ap4e1, Rab27b, Tmod2, Nes* and *Mylip,* and longer bout lengths were found in *Ghrhr* (Figure 3).

**Figure 3:**
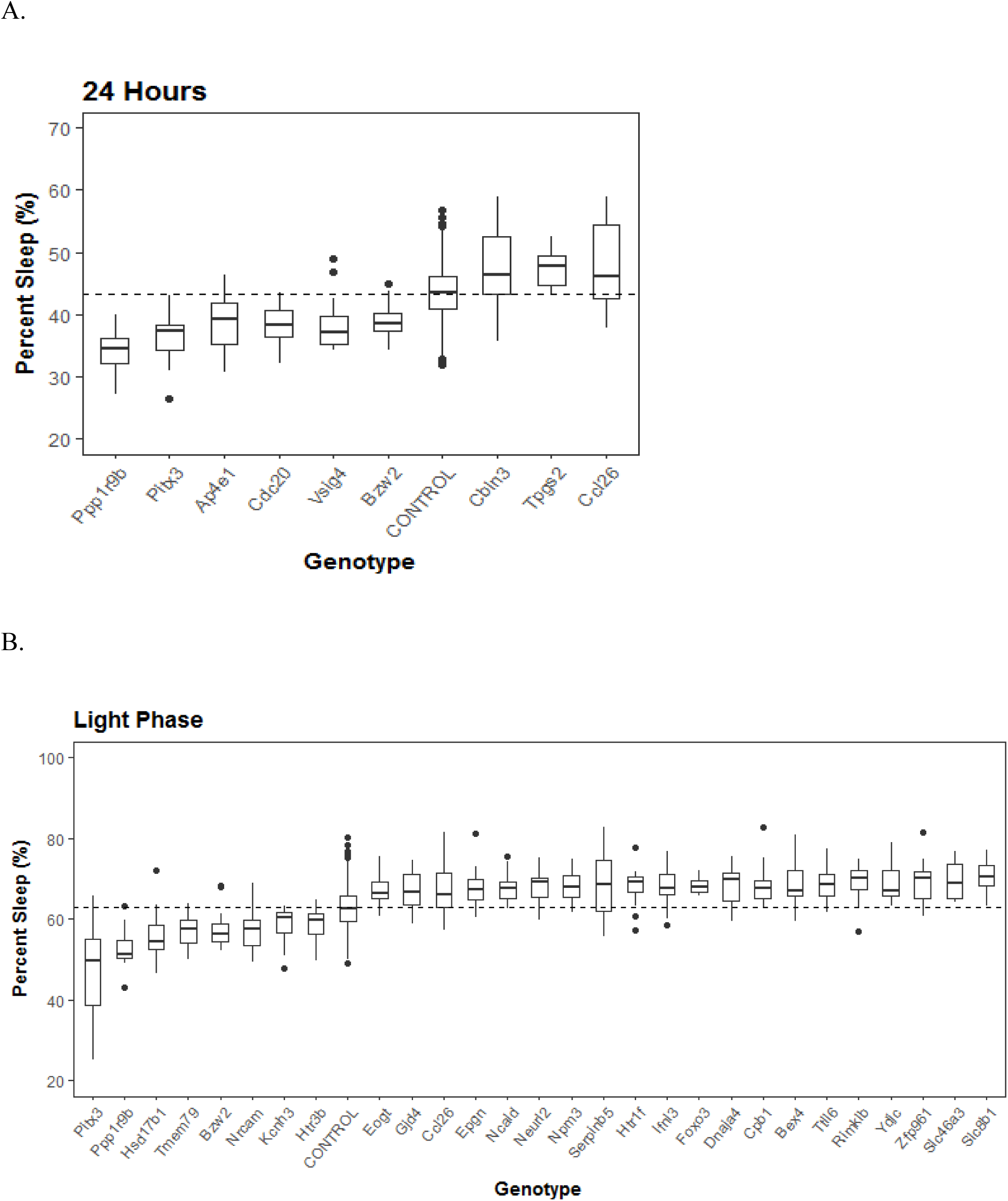
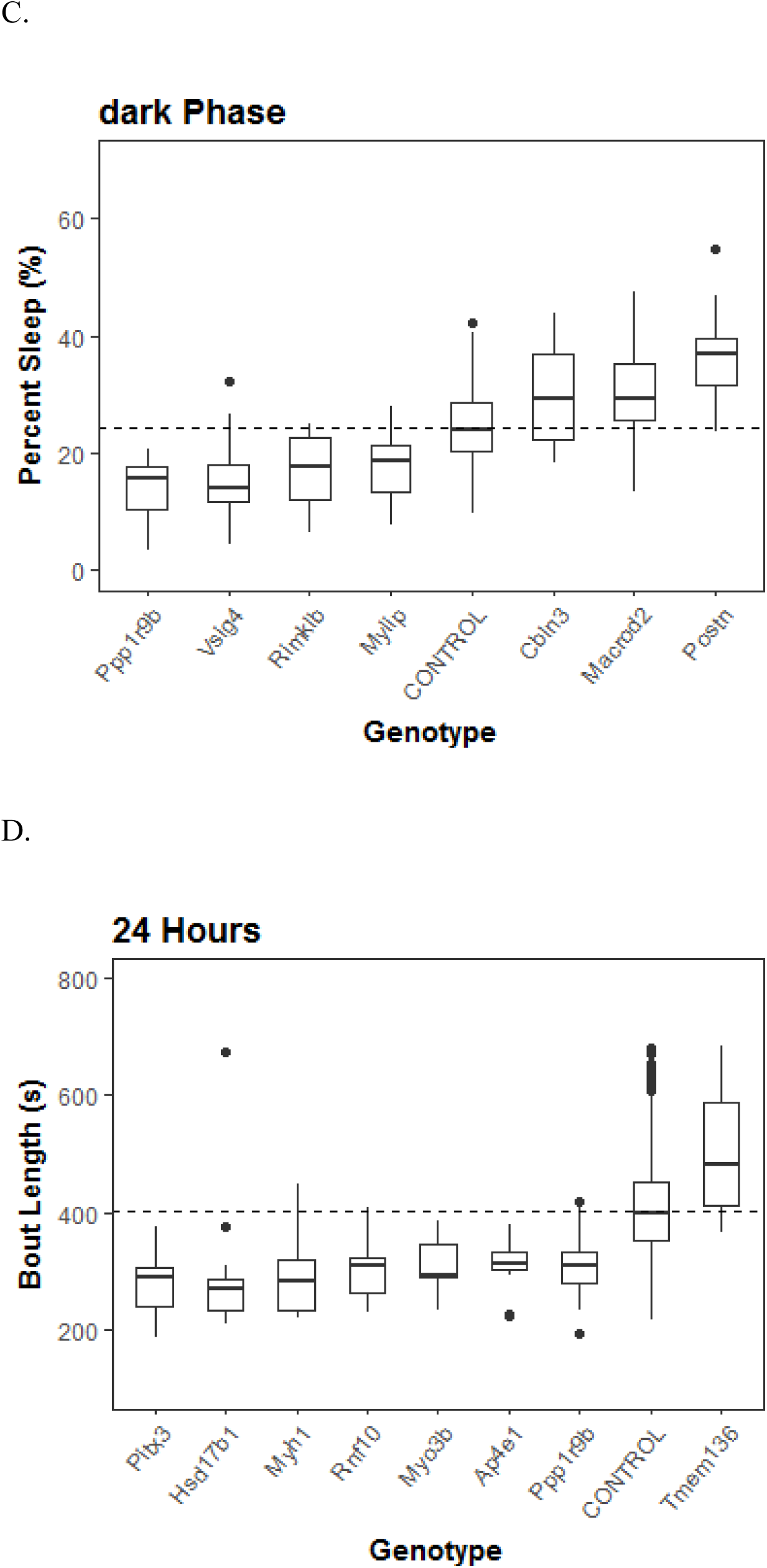
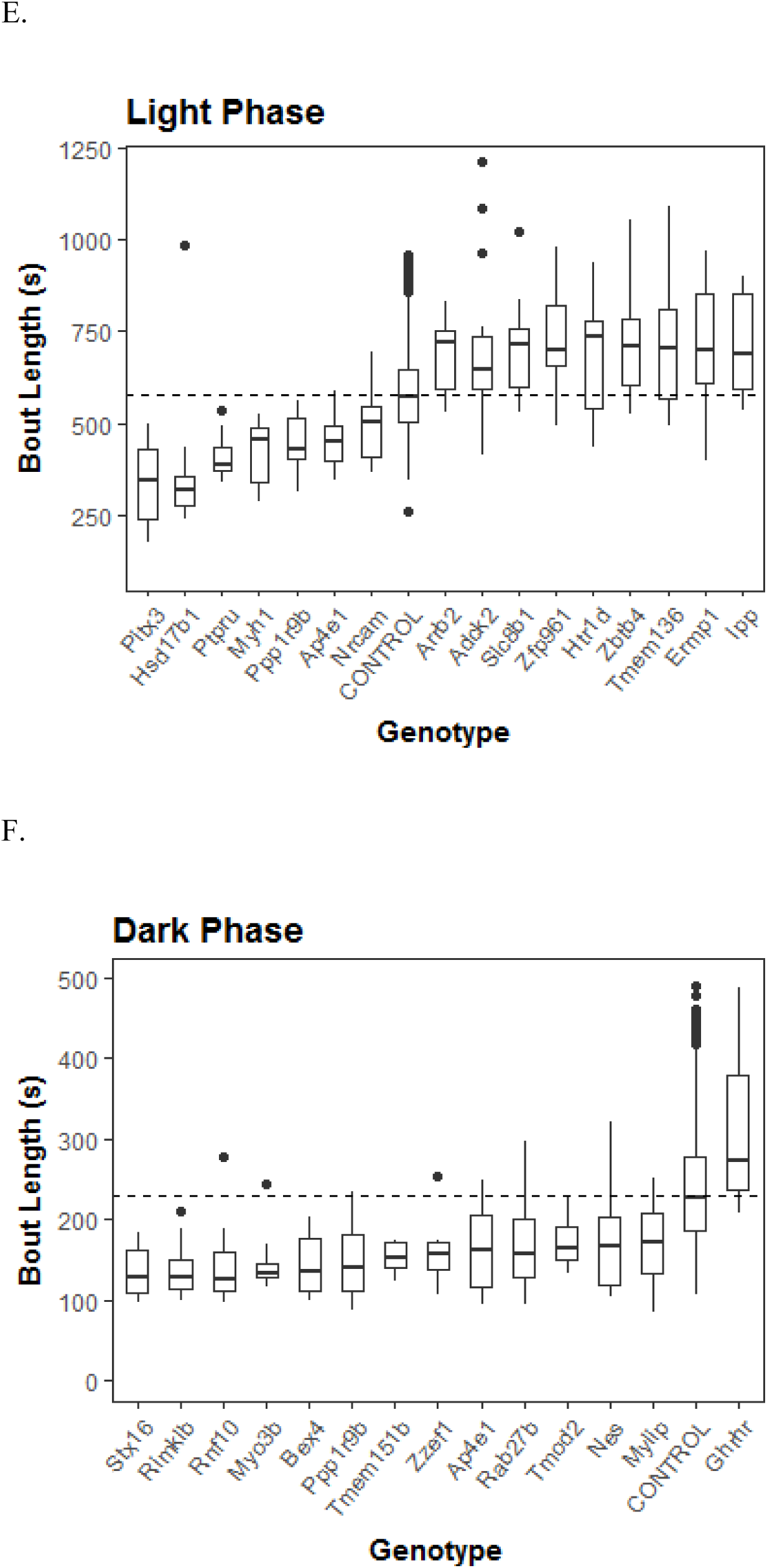
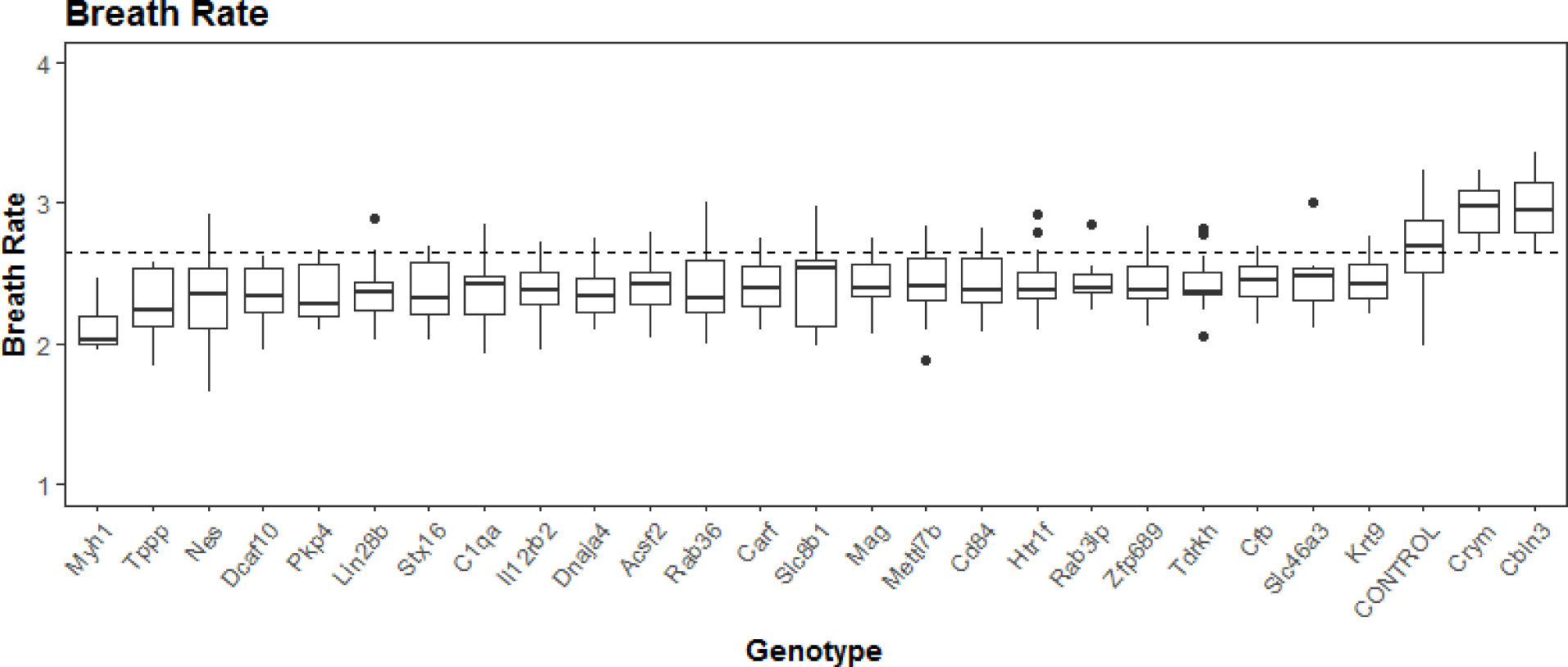
Knockout strains that differed significantly from control mice in percent sleep across 24 h (A), in light phase (B) and in dark phase (C); and bout length (D, E, F) across respective phases. (G) KO strains that differed significantly compared to controls in their breath rates.

### Genes affecting specific sex

Similar analysis (as described above) was performed for 289 KO strains, each of which had data for at least 3 females and 3 males to evaluate sex-specific differences in sleep parameters measured. Effect sizes have been reported as eta squared (Ƞ^2^) which represents the variance in each variable explained by either genotype, sex, or an interaction between them. According to Cohen’s guidelines, effect sizes can be defined as small (0.01), medium (0.06), and large (0.13) (46). Along with significant simple effects of genotype and sex, a significant interaction effect was observed for sleep percent (p<0.05; Ƞ^2^= 0.061) and mean bout lengths (p<0.06; Ƞ^2^= 0.053) (Table 1).

**Table 1:** Effect size and p-values for sex-genotype interactions in PiezoSleep variables.

| Piezo Variable | Effect Size ($\eta^2$) | p-value |
| --- | --- | --- |
| Daily Sleep Percent (24 h) | 0.061 | p<0.05 |
| Light Phase Sleep Percent | 0.053 | p<0.06 |
| Dark Phase Sleep Percent | 0.052 | p<0.07 |
| Bout Length mean (24 h) | 0.047 | p<0.08 |
| Light Phase Bout Length | 0.044 | p<0.09 |
| Dark Phase Bout Length | 0.051 | p<0.10 |
| Breath Rate during Sleep | 0.046 | p<0.11 |

Table 2 summarizes individual genes significant for sleep percent and bout length in one or both sexes and is graphically represented in Figure 4. For sleep percent over 24 hours, *Pitx3* and *Ppp1r9b* had significantly reduced sleep in both males and females. Sleep was significantly increased in females for *Cbln3, Ccl26, Rab24,* and *Ydjc*. During the light phase, both sexes had reduced sleep percent in *Pitx3*, and *Ppp1r9b*, and increased sleep percent in *Slc8b1*. In the dark phase, sleep durations were significantly reduced in *Ppp1r9b*, and increased in *Macrod2* and *Postn* in both sexes. Males had reduced sleep percent across 24 hrs, and dark phase, in *Mettl7b* and *Ptpn5,* and reduced dark phase percent in *Tmod.* Males had reduced bout length for *Mettl7b* and females had reduced bout length for *Ptpru*. Bout length was increased in males of *Nfatc4* and *Nat1*, and increased in females for *Slc8b1, Adck2, Zbtb4, Ipp,* and *Nrn1l*.

**Table 2:**
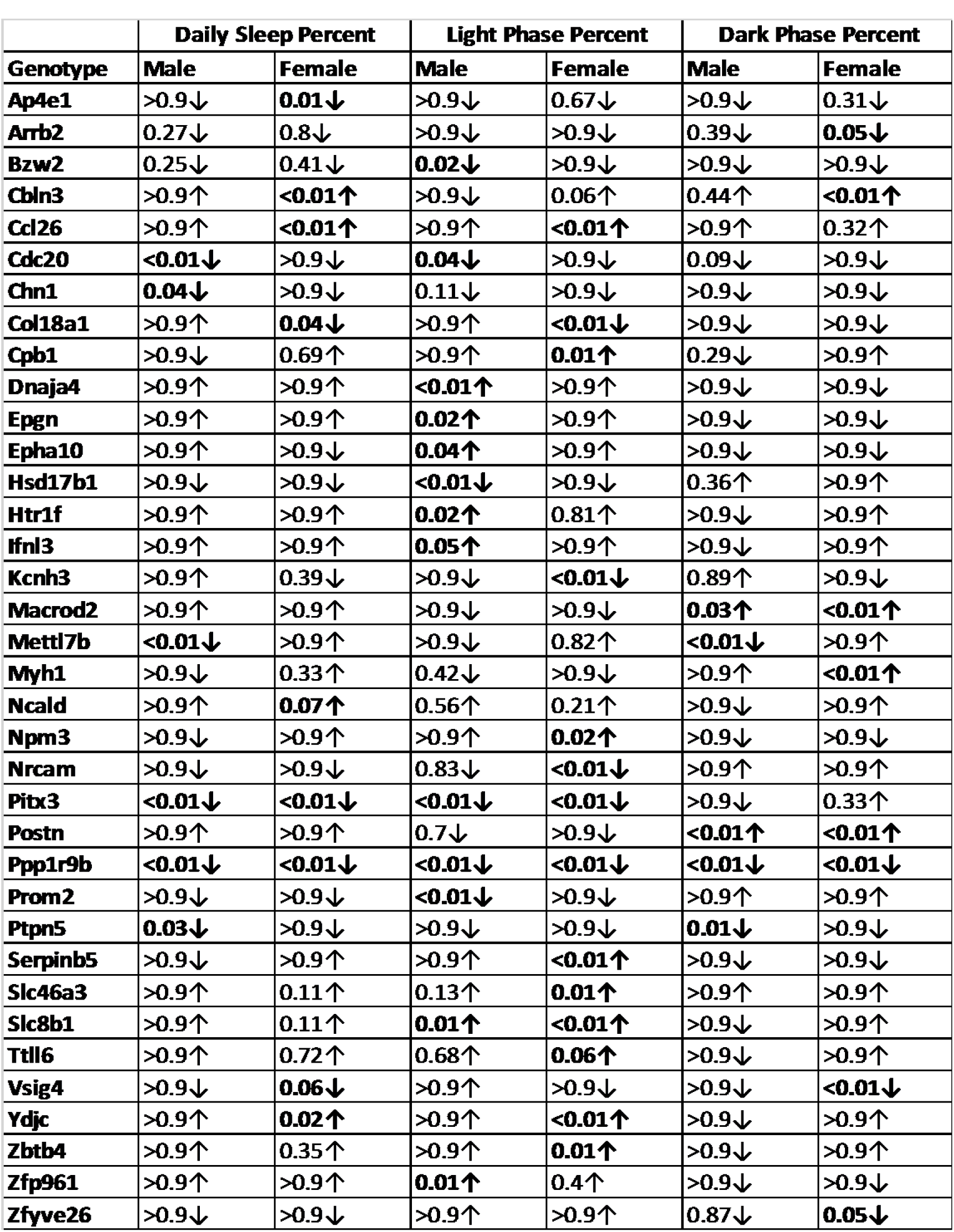
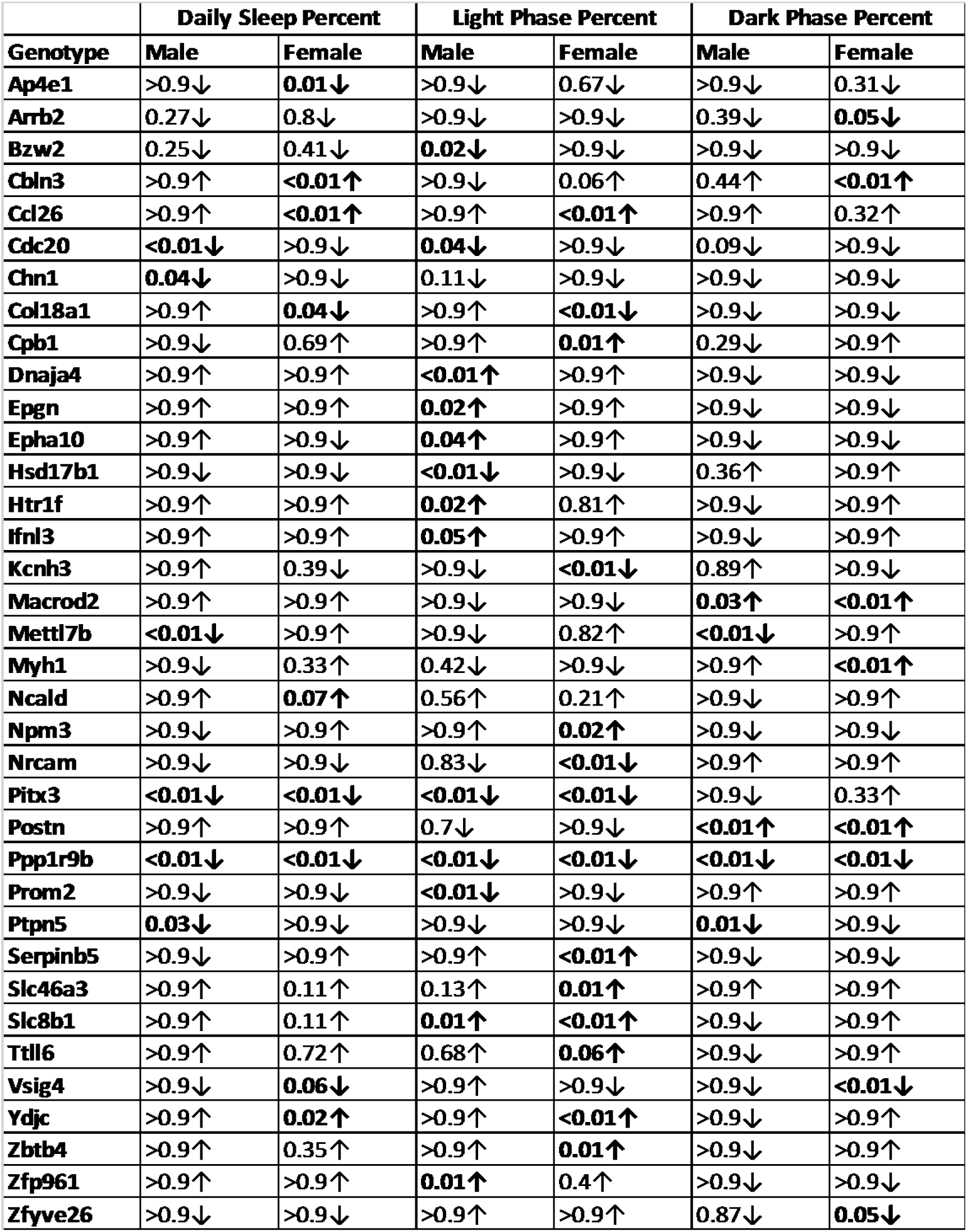
Sex differences in genotypes for (A) sleep percent and (B) bout lengths with direction of change. ↑ represents values higher than control animals, and ↓ represent values lower than controls. Bold arrows represent significant results.

|  | Daily Sleep Percent |  | Light Phase Percent |  | Dark Phase Percent |  |
| --- | --- | --- | --- | --- | --- | --- |
| Genotype | Male | Female | Male | Female | Male | Female |
| Ap4e1 | >0.9↓ | **0.01↓** | >0.9↓ | 0.67↓ | >0.9↓ | 0.31↓ |
| Arrb2 | 0.27↓ | 0.8↓ | >0.9↓ | >0.9↓ | 0.39↓ | **0.05↓** |
| Bzw2 | 0.25↓ | 0.41↓ | **0.02↓** | >0.9↓ | >0.9↓ | >0.9↓ |
| Cbln3 | >0.9↑ | **<0.01↑** | >0.9↓ | 0.06↑ | 0.44↑ | **<0.01↑** |
| Ccl26 | >0.9↑ | **<0.01↑** | >0.9↑ | **<0.01↑** | >0.9↑ | 0.32↑ |
| Cdc20 | **<0.01↓** | >0.9↓ | **0.04↓** | >0.9↓ | 0.09↓ | >0.9↓ |
| Chn1 | **0.04↓** | >0.9↓ | 0.11↓ | >0.9↓ | >0.9↓ | >0.9↓ |
| Col18a1 | >0.9↑ | **0.04↓** | >0.9↑ | **<0.01↓** | >0.9↓ | >0.9↓ |
| Cpb1 | >0.9↓ | 0.69↑ | >0.9↑ | **0.01↑** | 0.29↓ | >0.9↑ |
| Dnaja4 | >0.9↑ | >0.9↑ | **<0.01↑** | >0.9↑ | >0.9↓ | >0.9↓ |
| Epgn | >0.9↑ | >0.9↑ | **0.02↑** | >0.9↑ | >0.9↓ | >0.9↓ |
| Epha10 | >0.9↑ | >0.9↓ | **0.04↑** | >0.9↑ | >0.9↓ | >0.9↓ |
| Hsd17b1 | >0.9↓ | >0.9↓ | **<0.01↓** | >0.9↓ | 0.36↑ | >0.9↑ |
| Htr1f | >0.9↑ | >0.9↑ | **0.02↑** | 0.81↑ | >0.9↓ | >0.9↑ |
| Ifnl3 | >0.9↑ | >0.9↑ | **0.05↑** | >0.9↑ | >0.9↓ | >0.9↑ |
| Kcnh3 | >0.9↑ | 0.39↓ | >0.9↓ | **<0.01↓** | 0.89↑ | >0.9↓ |
| MacroD2 | >0.9↑ | >0.9↑ | >0.9↓ | >0.9↓ | **0.03↑** | **<0.01↑** |
| Mettl7b | **<0.01↓** | >0.9↑ | >0.9↓ | 0.82↑ | **<0.01↓** | >0.9↑ |
| Myh1 | >0.9↓ | 0.33↑ | 0.42↓ | >0.9↓ | >0.9↑ | **<0.01↑** |
| Ncald | >0.9↑ | **0.07↑** | 0.56↑ | 0.21↑ | >0.9↓ | >0.9↑ |
| Npm3 | >0.9↓ | >0.9↑ | >0.9↑ | **0.02↑** | >0.9↓ | >0.9↓ |
| Nrcam | >0.9↓ | >0.9↓ | 0.83↓ | **<0.01↓** | >0.9↑ | >0.9↑ |
| Pitx3 | **<0.01↓** | **<0.01↓** | **<0.01↓** | **<0.01↓** | >0.9↓ | 0.33↑ |
| Postn | >0.9↑ | >0.9↑ | 0.7↓ | >0.9↓ | **<0.01↑** | **<0.01↑** |
| Ppp1r9b | **<0.01↓** | **<0.01↓** | **<0.01↓** | **<0.01↓** | **<0.01↓** | **<0.01↓** |
| Prom2 | >0.9↓ | >0.9↓ | **<0.01↓** | >0.9↓ | >0.9↑ | >0.9↑ |
| Ptpn5 | **0.03↓** | >0.9↓ | >0.9↓ | >0.9↓ | **0.01↓** | >0.9↓ |
| Serpinb5 | >0.9↓ | >0.9↑ | >0.9↑ | **<0.01↑** | >0.9↓ | >0.9↓ |
| Slc46a3 | >0.9↑ | 0.11↑ | 0.13↑ | **0.01↑** | >0.9↑ | >0.9↑ |
| Slc8b1 | >0.9↑ | 0.11↑ | **0.01↑** | **<0.01↑** | >0.9↓ | >0.9↑ |
| Ttll6 | >0.9↑ | 0.72↑ | 0.68↑ | **0.06↑** | >0.9↓ | >0.9↑ |
| Vsig4 | >0.9↓ | **0.06↓** | >0.9↑ | >0.9↓ | >0.9↓ | **<0.01↓** |
| Ydjc | >0.9↑ | **0.02↑** | >0.9↑ | **<0.01↑** | >0.9↓ | >0.9↑ |
| Zbtb4 | >0.9↑ | 0.35↑ | >0.9↑ | **0.01↑** | >0.9↓ | >0.9↑ |
| Zfp961 | >0.9↑ | >0.9↑ | **0.01↑** | 0.4↑ | >0.9↓ | >0.9↓ |
| Zfyve26 | >0.9↓ | >0.9↓ | >0.9↑ | >0.9↑ | 0.87↓ | **0.05↓** |

**Figure 4:**
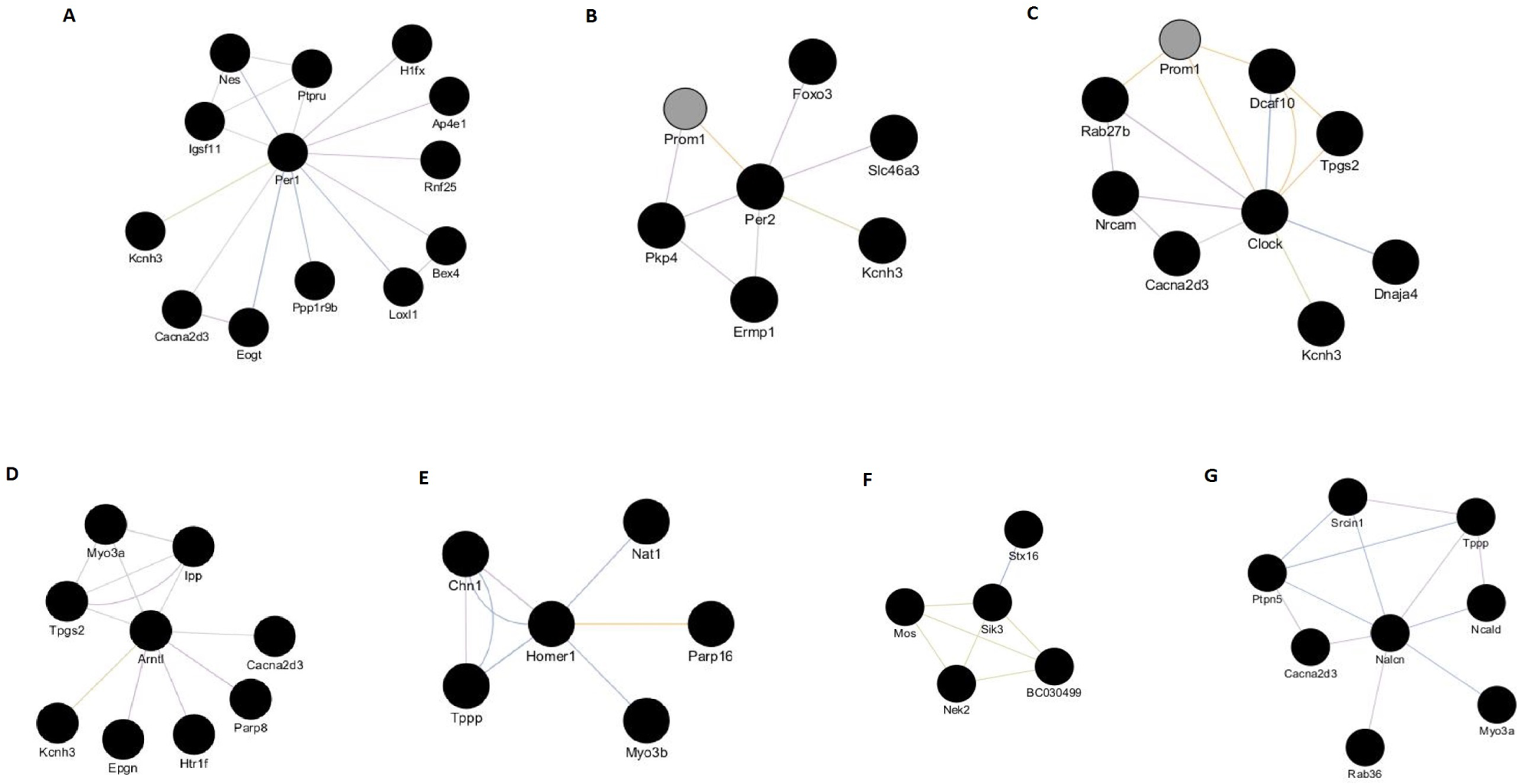
Gene networks predicted by GeneMANIA. Gene networks were created for previously known circadian and sleep related genes: A. Per1, B. Per2, C. Clock, D. Bmal1/Arntl, E. Homer1, F. Sik3, G. Nalcn. These networks are based on genetic interactions in which each node represents a single gene, and the edges represent physical associations, shared protein domains, co-expression, co-localization and their participation in the shared pathways for the candidate genes of interest.

### Principal Component Analysis

We performed additional analyses to identify sleep related genes based on multiple measures using Principal Component Analysis (PCA). We included standardized values for sleep percent and bout lengths for 24 hours, light phase and dark phase in our analysis. For our dataset, the first three principal components explained more than 95% of variability in the data, with PC1 accounting for 46.75% of the variability (Supplementary Figure 3). Correlation between the principal component and the original variables is described in terms of loadings (eigenvectors) (Supplementary Table 1). We calculated that 3 principal components best represent the data. Biplot reveals the relationship between different sleep variables in the first two principal components.

PC1 had high loading values for each variable (mean loading = 0.68; Figure 5C). ANOVA with multiple comparison through Dunnett’s post hoc test of PC1 resulted in 7 genotypes that showed significance. These are *Ap4e1, Cdc20, Hsd17b1, Myo3b, Pitx3, Ppp1r9b,* and *Rnf10*. Interestingly, PC2 had positive loadings for sleep percent and negative loadings for bout lengths, and *Arrb2, Cbln3, Dcaf10, Macrod2, Myh1, Parp8, Ppp1r9b, Rnf25, Tmem79,* and *Tpgs2* were significant. Similarly, PC3 had positive loadings for light phase and negative loadings for dark phase. The genes significant for PC3 include *Adck2, Ajap1, Arf2, Bex4, Cldn13, Cpb1, Dnaja4, Dnajc14, Eogt, Epgn, Epha10, Gipc3, H1fx, Hsd17b1, Htr1f, Ifnl3, Ipp, Kcnh3, Macrod2, Mylip, Nfatc4, Npm3, Nrcam, Parp16, Pitx3, Postn, Rab3, Rimklb, Serpinb5, Slc1a1, Slc46a3, Slc8b1, Stx16, Tdrkh, Tmod2, Ttll6, Zfp219,* and *Zfp961*. Based on the results from ANOVA of individual variables and PC3, we propose these genes as candidate genes that affect sleep in a specific circadian phase.

### Outlier Analysis

The analysis through ANOVA and PCA is based on the differences between knockout strains and controls and identifies strains that deviate from controls. As an additional strategy to identify genes affecting sleep, Mahalanobis distances (MD) for multivariate outliers were calculated for means of sleep percent and bout length variables of all KO strains. Candidate genes were identified with MD above a cutoff value of 3.80 (Chi-square alpha of 0.025). The advantage of this analysis is that it helps identify strains with consistently extreme values on multiple measures that might be missed while comparing single measures between mutants and controls. Genes that were significant earlier and appeared again in this analysis, provided additional validation of the results. Based on multivariate outlier analysis, the top candidate genes for sleep effects are *Akr1d1, Cacna2d3, Enox1, Fndc4, Galnt12, Gpr156, Hsd17b1, Ipp, Masp1, Mos, Myh1, Myo3a, Nek2, Ovch2, Pfdn6, Pitx3, Postn, Ppp1r9b, Ptpru, Rab3c, Rab3gap2, Rbm4b, Serpinb5, Srcin1, Tas2r138, Tmem79, Tnfsf18, Tubb4a,* and *Vsig4*.

### Key Candidate Genes for Sleep

The significant gene lists from ANOVA, PCA and MD analysis were compared to identify genes that appear in all of them. We consider these to be key candidate genes affecting sleep percent and bout lengths over 24 hours and light and dark phases. These genes are *Hsd17b1, Ipp, Myh1, Pitx3, Postn, Ppp1r9b, Serpinb5,* and *Tmem79* (Supplementary Figure 3). A more complete list of all 122 sleep related candidate genes was created by combining results from all of our analyses. This gene list was used for circadian and sleep gene association analysis, associations with IMPC phenotypes, and to find similar gene sets from the GeneWeaver database to perform preliminary functional annotations with respect to sleep.

### Associations with previously known sleep and circadian genes

GeneMANIA is a tool to identify associations between genes from large sets of data that include protein and genetic interactions, pathways, co-expression, co-localization, protein domain similarity, and predicted networks by OPHID (Online Predicted Human Interaction Database). We looked for associations with previously known circadian and sleep related genes. The genes selected for this analysis are known circadian clock genes showing differential expression during sleep and wake, and sleep alterations in mutant or knockout mice (*Clock, Bmal, Per1,* and *Per2*), and *Homer1* which is associated with homeostatic regulation of sleep. Two recently reported genes *Sik3* and *Nalcn* that regulate or influence non-rapid eye movement (NREM) and rapid eye movement (REM) sleep (Kunato-Yanagisawa 2016) were also included in this analysis. Associations were represented in the network format for each of the aforementioned genes in combination with our candidate genes. Each of our candidate genes was assessed for the number of associations with circadian and sleep related genes. *Kcnh3* was found to be associated with all four circadian genes. *Cacna2d3* had associations with *Per1, Clock, Bmal1,* and *Nalcn*. Two associations were observed for *Prom1* (with *Per2* and *Clock*), *Tpgs2* (with *Clock* and *Bmal1/Arntl*), and *Myo3a* (with *Bmal1/Arntl* and *Nalcn*) (Figure 4, Supplementary Table 2).

### Breath Rate

Breath rate during sleep was analyzed using the same approach used for other PiezoSleep parameters. We found 24 genotypes with significantly lower breath rate and two with higher breath rate (Figure 4G). Among these, *Acsf2, Lin28b, Myh1, Nes, Rab36,* and *Tppp* were significant for both females and males. Genes with male-specific effects include *Mettl7b, Ppp1r9b, Epgn, Postn, Stx16, Tmem151b, Zzef1, Adck2, Ajap1, BC030499, C1qa, Carf, Cers5, Ces4a, Cfb, Dcaf10, Il12rb2, Loxl1, Mag, Pkp4, Rab3ip, Rxfp4, Slc1a1, Tdrkh, Thsd1,* and *Zfp689* in which breath rate was significantly lower and *Prokr1* in which breath rate was higher. In females, breath rate was significantly higher than controls for *Cbln3, Crym*, and *Igsf11*.

### Coincident phenotype associations for top candidate genes

Abundant pleiotropy has been reported for complex traits (47). Given that sleep is a complex trait and almost a third of the genes in our analysis affect sleep, pleiotropy needs to be taken into account, especially given the likelihood that no “sleep-specific” genes exist. With a multitude of phenotypes and rich data in the KOMP2 pipeline, we performed cross trait analysis to identify coincident phenotypes for our top candidate genes. We analyzed the IMPC database for gene-phenotype associations, where phenotypes are reported as mammalian phenotype (MP) ontology terms. Our candidate genotypes resulted in 550 associations with 146 unique MP terms. Among the MP terms that showed high associations with genes, 37 had “abnormal sleep behavior”, 23 had “abnormal behavioral response to light”, 22 had “decreased circulating glucose level”, 20 had “decreased total body fat amount”, 16 had “abnormal behavior”, and 14 had associations to “hyperactivity”, “abnormal bone structure”, and “decreased bone mineral content”. When considered in terms of phenotype ontologies, more than half of the 550 associations observed belonged to either behavioral/neurological phenotype (167 associations) or homeostasis/metabolism phenotype (117 associations). These were followed by hematopoietic system phenotype with 65, growth/size/body region phenotype with 45, skeleton phenotype with 40, cardiovascular system phenotype with 30, adipose tissue phenotype with 20, and vision/eye phenotype with 16 associations (Supplementary Figures 5 and 6, Supplementary Table 3).

### Functional annotation of candidate genes

We used GeneWeaver to compare our candidate genes with pre-existing functional genomic data. GeneWeaver contains a database and a suite of tools that enable cross species and cross platform comparison of data from multiple genomic experiments. It provides output in the form of gene lists and computational tools to perform analysis that help in identifying genes that have been implicated previously in the phenotype of user’s interest. By entering the keyword “sleep” as the search term, we identified 26 gene sets of interest from mouse and human. Using Jaccard Similarity Tool for pairwise comparisons, 10 gene sets were identified that contained one or more of our key candidate genes (Supplementary Figure 7). Six datasets that are “Sleep Disorders”, “Sleep Stages”, “Sleep Deprivation”, “Abnormal Sleep Pattern”, “REM Sleep”, and “Abnormal frequency of paradoxical sleep” contained only one gene, the growth hormone releasing hormone receptor (*Ghrhr*). The other four genesets were QTL data for Dps (Delta power in slow wave sleep), and Rapid Eye Movement (REM) Sleep. *Tmem151b, Tubb4a, Cfb,* and *Pfdn6* were in the QTL region for Dps3 on chromosome 16, and *Tppp* in Dps1 QTL on chromosome 13. *Galnt12* and *Dcaf10* were in RemSlp1 QTL on chromosome 4, and *Htr1f, Igsf11,* and *Gpr156* in RemSlp3 QTL on chromosome 16.

## DISCUSSION

We report here the results from a large-scale phenotyping study that systematically assesses sleep in knockout mice. With sleep-wake recordings from more than 6000 mice, consisting of more than 1800 controls and 4500 mice representing 343 gene knockout strains, the KOMP2 pipeline at The Jackson Laboratory has generated a wealth of information-rich gene centric data that is unprecedented in sleep research. Through our analysis, we have identified genes previously not known to influence sleep. An experimental design using both sexes and the PiezoSleep monitoring system defining multiple sleep variables has allowed an in-depth analysis to identify genes that affect sleep in one or both sexes and during specific times of day. We found a remarkably high hit rate with approximately one third (35%) of gene knockouts tested identified with some degree of disordered sleep. This is perhaps not surprising given the dramatic changes in brain physiology and function that occur with sleep and wake transitions, and further suggests a potential to uncover many as yet unappreciated pathways affecting sleep. Although the purpose and functions of sleep are still unclear, we know that sleep is an important (and often essential) physiological process, and that even modest reductions of sleep in humans have substantial effects on health and cognitive functions (48–51).

We identified several genes that had significant effect on sleep only during a specific circadian or diurnal phase (i.e. light period vs. dark period). *Pitx3* (paired-like homeodomain transcription factor 3), *Bzw2* (basic leucine zipper W2 domains 2), and *Ccl26* (chemokine (C-C motif) ligand 26) affect sleep durations only during the light phase, while *Vsig4* (V-set and immunoglobulin domain containing 4) and *Cbln3* (cerebellin 3) do so only during the dark phase. Similarly, *Pitx3, Myh1* (myosin heavy polypeptide 1), and *Tmem136* (transmembrane protein 136) affect bout lengths only during the light phase, while *Myo3b* (myosin IIIB), and *Rnf10* (ring finger protein 10) affect bout lengths only during the dark phase. Most notable among these is the *Pitx3* gene knockout with markedly reduced sleep percent and bout length only during the light phase. Although little is known about the role of *Pitx3* in sleep, *Pitx3* is well known for its role in regulating lens development and is therefore associated with ocular abnormalities as seen in a range of animals including xenopus, zebrafish, humans as well as mice where its deficiency is reflected as a form of aphakia (52–54). A case study by Derwinska et al has reported that a hemizygous deletion on chromosome 10 including *Pitx3* resulted in sleep disturbances beginning in early childhood in a Caucasian boy (55). In the KOMP2 pipeline, *Pitx3* is associated with a multitude of additional outlier phenotypes including vision/eye, neurological/behavior, growth/size, homeostasis/metabolism, cardiovascular and skeletal assessments. In accordance with its known genetic function, *Pitx3* KO mice have anophthalmia or absence of eyes. In mice, *Pitx3* is also thought to be essential in development of dopaminergic neurons in Substantia Nigra (SN). Besides SN, *Pitx3* is also expressed in the Ventral Tegmental area (VTA) (56). Not only are these regions associated with reward, addiction and movement, they also play an important role in sleep and alertness (57). Interestingly, in 16 sleep-related candidate genes that we identified, including *Pitx3*, *Cbln3*, *Hsd17b1*, and *Rnf10*, we observed a vision/eye phenotype, making it important to further investigate the role of eye morphology and function in maintenance of sleep architecture. Note, however, that simple blindness or loss of eyesight entirely does not produce these same phenotypes (58–60).

Along with *Pitx3*, *Ppp1r9b* and *Ap4e1* are the candidate genes that were found to be most significant for both sleep percent and bout length. *Ppp1r9b* affects both sleep percent and bout lengths over 24 hours, light phase and dark phase. *Ppp1r9b* encodes the protein phosphatase 1 regulatory subunit 9B (also called neurabin II or spinophilin), and as the name indicates is a regulatory subunit of protein phosphatase (PP1). The protein product is highly enriched in dendritic spines (61). Sleep is reported to promote formation of dendritic spines for memory consolidation (62). In addition, PP1 regulates AMPA channels that are believed to play a role in synaptic plasticity, and learning and memory (63). Further, *PPP1R9B* is one of the substrates of GSK3β (glycogen synthase kinase 3 beta) which is a crucial circadian clock regulator (64). Casein kinase I enzymes have been shown to play a critical role in regulating clock genes such as *Per2*, and *PPP1R9B* may dephosphorylate some of these same sites and work in opposition (65). There is increasing evidence that clock genes not only influence circadian aspects of sleep and wake, but are fundamentally tied to sleep homeostasis as well, which appears to be altered in the *Ppp1r9b* knockout mice (64,66). *Ap4e1* was found to be significant for 24 hours, light phase and dark phase for bout length, and sleep percent over 24 hours. It codes for the epsilon subunit 1 of adaptor protein (AP) 4 complex that is involved in vesicle trafficking. *Ap4e1* has been associated with cerebral palsy, and mutations in humans are known to cause intellectual disabilities, with abnormal sleep behavior reported as a coincident phenotype (67). Like *Pitx3*, *Ap4e1* is associated with multiple additional aberrant phenotypes observed in the KOMP2 pipeline, and is within the QTL Cplaq15 (circadian period of locomotor activity 15) (68).

Another KO of interest is *Kcnh3 (Kv12.2)* that was significant for sleep percent in females, and was also found to be associated with previously known circadian pathway genes. It is a subunit of potassium channels that regulate neuronal excitability. *Kcnh3* KO mice have shorter sleep duration in the light phase, and was found to be associated with *Per1, Per2, Clock* and *Bmal1* through co-expression. A similar but less pronounced reduction in sleep duration was also seen in *Kcnh3* heterozygous mice (data not included). Its overexpression has been associated with deficits in learning, and its ablation with enhanced cognitive functions (spatial and working memory), hippocampal hyperexcitability and spontaneous seizures (69,70). Many other potassium channels are known to modulate sleep-wake. A well-known example is *Shaker* in drosophila (71). Flies mutant for the *Shaker* gene are short sleepers. In previous research, NREM sleep has been reported to be reduced in *Kcnc1, Kcnc3, Kcnc1/3 and Kv1.2* KO mice in magnitudes similar to what we have reported for Kcnh3 (8,10). Altered potassium channels may reduce the resting membrane potential of neurons, leading to increased firing and reduced sleep. While this would occur in both inhibitory and excitatory circuits, the net effect is presumably increased excitation. Evidence supporting this hypothesis was found in a recent study showing that sleep and anesthesia reduced extracellular potassium ion levels in the brain (72).

Several genes were found to affect breath rate, many of which did not have coincident effects on sleep, such as *Tppp* (p25 alpha/p24) mice which had a shorter breath rate. *Tppp* encodes tubulin-binding protein believed to be important for oligodendrocyte differentiation (73). So far it is not known if breath rate-regulating mechanisms are in any way associated with other brain functions or sleep-related pathways, but clear variations in breathing rhythms during sleep and wake suggests an underlying relationship between these processes (74). In addition, during the sleep phase, there are variations in breathing rate during REM and NREM stages. In the future, as algorithms to record and detect breathing rhythm changes in the PiezoSleep system are further developed, it will become possible to identify genes related to these sleep stages as well. From our analysis of geneset similarity with REM sleep QTLs and interactions within networks for *Sik3* and *Nalcn*, we propose 16 putative candidates for REM/NREM sleep. *Cacna2d3*, a calcium channel protein, is an interesting candidate since it interacts with *Nalcn* along with circadian *Per1*, *Clock* and *Bmal1* genes. Five of these genes including *Stx16, Tppp, Rab36, Dcaf10,* and *Htr1f* also significantly affect breath rate during sleep and should be a focus of further study. Based on similar evidence from gene network of *Homer1* and *Dps* QTLs, we propose 9 novel candidate genes for homeostatic regulation of sleep. Here again, *Tppp* is a key candidate gene since it interacts with *Homer1* in co-expression network, and is also a part of *Dps1* QTL.

Sexual dimorphism, although often overlooked, is prevalent in KOMP^2^ strains and controls for many phenotypes and was also observed in the investigated sleep parameters (75). In B6J mice, this dimorphism has also been reported for REM and NREM sleep stages, with lower values observed in female mice (76,77). Sex differences observed in human studies have been evident but generally smaller in magnitude (78,79). Sex hormones are thought to be one of the contributing factors for these differences (80–82). In our analysis, in addition to significant sex differences in control mice, the majority of our findings are genotypes that had highly significant sex specific differences in sleep traits. Further study of these genes might prove helpful in identifying causes for this intriguing sexual dimorphism seen in sleep behavior.

Although we detected many novel sleep genes, the extensive filtering of sleep signals used in our study most likely excluded genes affecting sleep in subtle ways. Genes involved in sleep that have extensive redundancy or compensatory mechanisms in place to mask the effect of a gene ablation would also be missed. Finally, the knockout method is in general limited by the fact that a gene is ablated in all tissues, so KO of essential genes that may in fact be involved in sleep result in a non-viable animal, preventing sleep phenotyping. However, utilization of the conditional allele obtained from these KOs may overcome this limitation. It is also unclear to what extent any of the sleep alterations from gene ablation are due to direct or indirect effects of the gene in question. These questions may be partially addressed by examining the multiple phenotypes for each knockout, which we have done in a rather limited way. There is no reason to believe that there are any genes whose sole functions are related to sleep, as even the so-called core circadian clock genes are pleiotropic.

Our study demonstrates the utility of rapid-non-invasive sleep phenotyping in high-throughput mouse screens. This initial set of approximately 350 genes was not selected to have sleep phenotypes and yet a high percentage were found to have altered sleep phenotypes, and of a magnitude as large as any that have been selected specifically for sleep studies over the past 25 years (10). This supports the utility of an unbiased selection and phenotyping for mouse knockouts, especially given that a majority of genes are not well understood. Unlike most individual studies of KO mice that examine genes predicted to influence a trait of interest, the IMPC/KOMP2 is a comprehensive, unbiased approach, having examined more than 5000 genes to date, and thus holds potential for detecting and identifying unexpected and pleiotropic effects of the knocked out genes (27). With fewer than 350 KOs analyzed for sleep to date, we demonstrate the potential of this large-scale effort to find novel sleep phenotypes for a significant percentage of coding genes. To the best of our knowledge, none of these genes have been previously implicated in sleep, further suggesting that most genes influencing sleep have yet to be identified.

## CONCLUSIONS

This study utilizes data generated from the KOMP2 project at The Jackson Laboratory, a large-scale project intended to phenotype knockout mice in alignment with the IMPC. We report on analysis of 6350 mice representing 343 different gene knockouts, along with over 1800 C57BL/6NJ control mice all assessed for sleep and wake as a part of the unique behavioral phenotyping pipeline at JAX. Our findings showed altered sleep and wake in many of the knockout lines compared to the controls. Many of these strains also exhibit sex differences in sleep traits. Among all the sleep related candidate genes that we have reported in this study, with the exception of *Ghrhr*, none have been implicated in sleep regulation to the best of our knowledge. *Pitx3* and *Ppp1r9b* would be the strongest sleep related candidate genes identified from our analysis that strongly impact sleep. Follow up studies and experiments for these and other candidate genes will potentially help in discovering new signaling pathways and gain better understanding of the many processes governing or influencing sleep. Our high hit rate, however, raises the issue that the genetic influences on sleep will involve a high percentage of the genome, and each individual mutation or knockout that influences sleep is perhaps unlikely to yield very much insight. This is true for our 3-5 biggest hits here and also the very interesting mutations found in the only other such large-scale sleep mutant screen (83,84). In that study, mutations of *Sik3* and *Nalcn* clearly impact sleep, but may simply be among hundreds or thousands that can alter sleep when mutated or deleted. Like most complex traits, the future may rely on systems genomics and systems biology to make sense of many interacting genes and pathways, as there may be no small set of genes that clearly perform any critical aspect of sleep functions or sleep regulation, other than perhaps the so-called “clock” genes that are central to the circadian pacemaker, and perhaps sleep homeostasis as well (64). Nonetheless, these newly identified genes that when perturbed alter sleep and may provide another early step towards understanding the mechanisms of sleep. Further study of these genes and members of their associated pathways may lead to enhanced understanding of the nature and role for sleep and may further provide novel targets for treatment of specific sleep disorders.

## ACKNOWLEDGEMENTS

This study was funded by NIH OD011185 and NIH HG006332.

We dedicate this paper to Martin Striz, who died unexpectedly in August 2014. He was an outstanding and dedicated student, teacher and scientist, and a kind and wonderful friend, who always gave generously of his time and expertise. He helped provide critical training and assistance to several individuals in this current work as well. He is missed daily, and will continue to be missed by all who knew him.

## References

1. Tucker MA, Hirota Y, Wamsley EJ, Lau H, Chaklader A, Fishbein W. A daytime nap containing solely non-REM sleep enhances declarative but not procedural memory. Neurobiol Learn Mem [Internet]. 2006;86(2):241–7. Available from: http://www.ncbi.nlm.nih.gov/pubmed/16647282

2. Diekelmann S, Born J. The memory function of sleep. Nat Rev Neurosci [Internet]. 2010;11(2):114–26. Available from: http://www.ncbi.nlm.nih.gov/pubmed/20046194

3. Marshall L, Born J. The contribution of sleep to hippocampus-dependent memory consolidation. Trends Cogn Sci [Internet]. 2007;11(10):442–50. Available from: http://www.ncbi.nlm.nih.gov/pubmed/17905642

4. Nishida M, Pearsall J, Buckner RL, Walker MP. REM sleep, prefrontal theta, and the consolidation of human emotional memory. Cereb Cortex [Internet]. 2009;19(5):1158–66. Available from: http://www.ncbi.nlm.nih.gov/pubmed/18832332

5. Xie L, Kang H, Xu Q, Chen MJ, Liao Y, Thiyagarajan M, et al. Sleep drives metabolite clearance from the adult brain. Science (80-) [Internet]. 2013;342(6156):373–7. Available from: http://www.ncbi.nlm.nih.gov/pubmed/24136970

6. Tononi G, Cirelli C. Sleep function and synaptic homeostasis. Sleep Med Rev [Internet]. 2006;10(1):49–62. Available from: http://www.ncbi.nlm.nih.gov/pubmed/16376591

7. Krueger JM, Frank MG, Wisor JP, Roy S. Sleep function: Toward elucidating an enigma. Sleep Med Rev [Internet]. 2016;28:46–54. Available from: http://www.ncbi.nlm.nih.gov/pubmed/26447948

8. Rechtschaffen A. Current perspectives on the function of sleep. Perspect Biol Med [Internet]. 1998;41(3):359–90. Available from: http://www.ncbi.nlm.nih.gov/pubmed/9604368

9. Vassalli A, Dijk DJ. Sleep function: current questions and new approaches. Eur J Neurosci [Internet]. 2009/05/29. 2009;29(9):1830–41. Available from: http://www.ncbi.nlm.nih.gov/pubmed/19473236

10. Cirelli C. The genetic and molecular regulation of sleep: from fruit flies to humans. Nat Rev Neurosci [Internet]. 2009;10(8):549–60. Available from: http://www.ncbi.nlm.nih.gov/pubmed/19617891

11. Gondo Y, Fukumura R, Murata T, Makino S. Next-generation gene targeting in the mouse for functional genomics. BMB Rep [Internet]. 2009;42(6):315–23. Available from: http://www.ncbi.nlm.nih.gov/pubmed/19558788

12. O’Hara BF, Ding J, Bernat RL, Franken P. Genomic and proteomic approaches towards an understanding of sleep. CNS Neurol Disord Drug Targets [Internet]. 2007;6(1):71–81. Available from: http://www.ncbi.nlm.nih.gov/pubmed/17305555

13. Vitaterna MH, King DP, Chang AM, Kornhauser JM, Lowrey PL, McDonald JD, et al. Mutagenesis and mapping of a mouse gene, Clock, essential for circadian behavior. Science (80-) [Internet]. 1994;264(5159):719–25. Available from: http://www.ncbi.nlm.nih.gov/pubmed/8171325

14. Kapfhamer D, Valladares O, Sun Y, Nolan PM, Rux JJ, Arnold SE, et al. Mutations in Rab3a alter circadian period and homeostatic response to sleep loss in the mouse. Nat Genet [Internet]. 2002/09/24. 2002;32(2):290–5. Available from: http://www.ncbi.nlm.nih.gov/pubmed/12244319

15. Kloss B, Price JL, Saez L, Blau J, Rothenfluh A, Wesley CS, et al. The Drosophila clock gene double-time encodes a protein closely related to human casein kinase Iepsilon. Cell [Internet]. 1998/07/23. 1998;94(1):97–107. Available from: http://www.ncbi.nlm.nih.gov/pubmed/9674431

16. Franken P, Chollet D, Tafti M. The homeostatic regulation of sleep need is under genetic control. J Neurosci [Internet]. 2001/04/18. 2001;21(8):2610–21. Available from: https://www.ncbi.nlm.nih.gov/pubmed/11306614

17. Nadeau JH. Modifier genes in mice and humans. Nat Rev Genet [Internet]. 2001;2(3):165–74. Available from: http://www.ncbi.nlm.nih.gov/entrez/query.fcgi?cmd=Retrieve&db=PubMed&dopt=Citation&list_uids=11256068

18. Tafti M, Petit B, Chollet D, Neidhart E, de Bilbao F, Kiss JZ, et al. Deficiency in short-chain fatty acid beta-oxidation affects theta oscillations during sleep. Nat Genet. 2003;34(3):320–5.

19. Drager UC. Retinoic acid signaling in the functioning brain. Sci STKE [Internet]. 2006;2006(324):pe10. Available from: http://www.ncbi.nlm.nih.gov/pubmed/16507818

20. Maret S, Dorsaz S, Gurcel L, Pradervand S, Petit B, Pfister C, et al. Homer1a is a core brain molecular correlate of sleep loss. Proc Natl Acad Sci U S A. 2007;104(50):20090–5.

21. O’Hara BF, Jiang P, Turek FW, Franken P. Chapter 29 - Genetics and Genomic Basis of Sleep in Rodents A2 - Kryger, Meir. In: Roth T, Dement WC, editors. Principles and Practice of Sleep Medicine (Sixth Edition) [Internet]. Elsevier; 2017. p. 296–309.e5. Available from: http://www.sciencedirect.com/science/article/pii/B9780323242882000295

22. Donohue KD, Medonza DC, Crane ER, O’Hara BF. Assessment of a non-invasive high-throughput classifier for behaviours associated with sleep and wake in mice. Biomed Eng Online. 2008;7:14.

23. Flores AE, Flores JE, Deshpande H, Picazo JA, Xie XMS, Franken P, et al. Pattern recognition of sleep in rodents using piezoelectric signals generated by gross body movements. Ieee Trans Biomed Eng. 2007;54(2):225–33.

24. Mang GM, Nicod J, Emmenegger Y, Donohue KD, O’Hara BF, Franken P. Evaluation of a piezoelectric system as an alternative to electroencephalogram/ electromyogram recordings in mouse sleep studies. Sleep [Internet]. 2014;37(8):1383–92. Available from: http://www.ncbi.nlm.nih.gov/pubmed/25083019

25. Philip VM, Sokoloff G, Ackert-Bicknell CL, Striz M, Branstetter L, Beckmann MA, et al. Genetic analysis in the Collaborative Cross breeding population. Genome Res [Internet]. 2011;21(8):1223–38. Available from: http://genome.cshlp.org/content/21/8/1223.abstract

26. Pack AI, Galante RJ, Maislin G, Cater J, Metaxas D, Lu S, et al. Novel method for high-throughput phenotyping of sleep in mice. Physiol Genomics [Internet]. 2007;28(2):232–8. Available from: http://www.ncbi.nlm.nih.gov/entrez/query.fcgi?cmd=Retrieve&db=PubMed&dopt=Citation&list_uids=16985007

27. Brown SD, Moore MW. Towards an encyclopaedia of mammalian gene function: the International Mouse Phenotyping Consortium. Dis Model Mech [Internet]. 2012/05/09. 2012;5(3):289–92. Available from: http://www.ncbi.nlm.nih.gov/pubmed/22566555

28. Ringwald M, Iyer V, Mason JC, Stone KR, Tadepally HD, Kadin JA, et al. The IKMC web portal: a central point of entry to data and resources from the International Knockout Mouse Consortium. Nucleic Acids Res [Internet]. 2010/10/12. 2011;39(Database issue):D849–55. Available from: http://www.ncbi.nlm.nih.gov/pubmed/20929875

29. Bradley A, Anastassiadis K, Ayadi A, Battey JF, Bell C, Birling MC, et al. The mammalian gene function resource: the International Knockout Mouse Consortium. Mamm Genome [Internet]. 2012/09/13. 2012;23(9–10):580–6. Available from: http://www.ncbi.nlm.nih.gov/pubmed/22968824

30. Abbott A. Mouse project to find each gene’s role. Nature [Internet]. 2010/05/28. 2010;465(7297):410. Available from: http://www.ncbi.nlm.nih.gov/pubmed/20505705

31. Brown SD, Moore MW. The International Mouse Phenotyping Consortium: past and future perspectives on mouse phenotyping. Mamm Genome [Internet]. 2012;23(9–10):632–40. Available from: http://www.ncbi.nlm.nih.gov/pubmed/22940749

32. Schofield PN, Hoehndorf R, Gkoutos G V. Mouse genetic and phenotypic resources for human genetics. Hum Mutat [Internet]. 2012;33(5):826–36. Available from: http://www.ncbi.nlm.nih.gov/pubmed/22422677

33. Skarnes WC, Rosen B, West AP, Koutsourakis M, Bushell W, Iyer V, et al. A conditional knockout resource for the genome-wide study of mouse gene function. Nature [Internet]. 2011;474(7351):337–42. Available from: http://www.ncbi.nlm.nih.gov/pubmed/21677750

34. Mekada K, Hirose M, Murakami A, Yoshiki A. Development of SNP markers for C57BL/6N-derived mouse inbred strains. Exp Anim [Internet]. 2014/10/25. 2015;64(1):91–100. Available from: http://www.ncbi.nlm.nih.gov/pubmed/25341966

35. Simon MM, Greenaway S, White JK, Fuchs H, Gailus-Durner V, Wells S, et al. A comparative phenotypic and genomic analysis of C57BL/6J and C57BL/6N mouse strains. Genome Biol [Internet]. 2013/08/02. 2013;14(7):R82. Available from: http://www.ncbi.nlm.nih.gov/pubmed/23902802

36. Keane TM, Goodstadt L, Danecek P, White MA, Wong K, Yalcin B, et al. Mouse genomic variation and its effect on phenotypes and gene regulation. Nature [Internet]. 2011/09/17. 2011;477(7364):289–94. Available from: http://www.ncbi.nlm.nih.gov/pubmed/21921910

37. Morgan H, Simon M, Mallon AM. Accessing and mining data from large-scale mouse phenotyping projects. Int Rev Neurobiol [Internet]. 2012;104:47–70. Available from: http://www.ncbi.nlm.nih.gov/pubmed/23195311

38. Mitchell AFS, Krzanowski WJ. The Mahalanobis Distance and Elliptic Distributions. Biometrika. 1985;72(2):464–7.

39. Bassett JH, Gogakos A, White JK, Evans H, Jacques RM, van der Spek AH, et al. Rapid-throughput skeletal phenotyping of 100 knockout mice identifies 9 new genes that determine bone strength. PLoS Genet [Internet]. 2012/08/10. 2012;8(8):e1002858. Available from: http://www.ncbi.nlm.nih.gov/pubmed/22876197

40. Warde-Farley D, Donaldson SL, Comes O, Zuberi K, Badrawi R, Chao P, et al. The GeneMANIA prediction server: biological network integration for gene prioritization and predicting gene function. Nucleic Acids Res [Internet]. 2010;38(Web Server issue):W214–20. Available from: http://www.ncbi.nlm.nih.gov/pubmed/20576703

41. Bubier JA, Langston MA, Baker EJ, Chesler EJ. Integrative Functional Genomics for Systems Genetics in GeneWeaver.org. Methods Mol Biol [Internet]. 2017;1488:131–52. Available from: https://www.ncbi.nlm.nih.gov/pubmed/27933523

42. R Development Core Team R. R: A Language and Environment for Statistical Computing. R Foundation for Statistical Computing. 2011.

43. Garcia H, Filzmoser P. Multivariate Statistical Analysis using the R package chemometrics. Vienna: Austria. 2011;

44. Signorell A, others. DescTools: Tools for descriptive statistics. R Packag version 099. 2016;18.

45. Wickham H. Ggplot2. Elegant Graphics for Data Analysis. 2009.

46. Cohen J. A power primer. Psychol Bull [Internet]. 1992;112(1):155–9. Available from: https://www.ncbi.nlm.nih.gov/pubmed/19565683

47. Sivakumaran S, Agakov F, Theodoratou E, Prendergast JG, Zgaga L, Manolio T, et al. Abundant pleiotropy in human complex diseases and traits. Am J Hum Genet [Internet]. 2011;89(5):607–18. Available from: https://www.ncbi.nlm.nih.gov/pubmed/22077970

48. Shaw PJ, Franken P. Perchance to dream: solving the mystery of sleep through genetic analysis. J Neurobiol [Internet]. 2002/12/18. 2003;54(1):179–202. Available from: http://www.ncbi.nlm.nih.gov/pubmed/12486704

49. Cohen S, Doyle WJ, Alper CM, Janicki-Deverts D, Turner RB. Sleep habits and susceptibility to the common cold. Arch Intern Med [Internet]. 2009/01/14. 2009;169(1):62–7. Available from: http://www.ncbi.nlm.nih.gov/pubmed/19139325

50. Durmer JS, Dinges DF. Neurocognitive consequences of sleep deprivation. Semin Neurol [Internet]. 2005/03/31. 2005;25(1):117–29. Available from: http://www.ncbi.nlm.nih.gov/pubmed/15798944

51. Cespedes EM, Bhupathiraju SN, Li Y, Rosner B, Redline S, Hu FB. Long-term changes in sleep duration, energy balance and risk of type 2 diabetes. Diabetologia [Internet]. 2016;59(1):101–9. Available from: http://www.ncbi.nlm.nih.gov/pubmed/26522276

52. Huang B, He W. Molecular characteristics of inherited congenital cataracts. Eur J Med Genet [Internet]. 2010;53(6):347–57. Available from: http://www.ncbi.nlm.nih.gov/pubmed/20624502

53. Khosrowshahian F, Wolanski M, Chang WY, Fujiki K, Jacobs L, Crawford MJ. Lens and retina formation require expression of Pitx3 in Xenopus pre-lens ectoderm. Dev Dyn [Internet]. 2005;234(3):577–89. Available from: http://www.ncbi.nlm.nih.gov/pubmed/16170783

54. Shi X, Bosenko D V, Zinkevich NS, Foley S, Hyde DR, Semina E V, et al. Zebrafish pitx3 is necessary for normal lens and retinal development. Mech Dev [Internet]. 2005;122(4):513–27. Available from: http://www.ncbi.nlm.nih.gov/pubmed/15804565

55. Derwinska K, Mierzewska H, Goszczanska A, Szczepanik E, Xia ZL, Kusmierska K, et al. Clinical improvement of the aggressive neurobehavioral phenotype in a patient with a deletion of PITX3 and the absence of L-DOPA in the cerebrospinal fluid. Am J Med Genet Part B-Neuropsychiatric Genet. 2012;159B(2):236–42.

56. Li J, Dani JA, Le W. The role of transcription factor Pitx3 in dopamine neuron development and Parkinson’s disease. Curr Top Med Chem [Internet]. 2009;9(10):855–9. Available from: http://www.ncbi.nlm.nih.gov/pubmed/19754401

57. Nishino S. Neurotransmitters and Neuropharmacology of Sleep/Wake Regulations A2 - Kushida, Clete A. In: Encyclopedia of Sleep [Internet]. Waltham: Academic Press; 2013. p. 395–406. Available from: http://www.sciencedirect.com/science/article/pii/B9780123786104000875

58. Lucas RJ, Freedman MS, Lupi D, Munoz M, David-Gray ZK, Foster RG. Identifying the photoreceptive inputs to the mammalian circadian system using transgenic and retinally degenerate mice. Behav Brain Res. 2001;

59. Freedman MS, Lucas RJ, Soni B, Von Schantz M, Muñoz M, David-Gray Z, et al. Regulation of mammalian circadian behavior by non-rod, non-cone, ocular photoreceptors. Science (80-). 1999;

60. Argamaso SM, Froehlich AC, McCall MA, Nevo E, Provencio I, Foster RG. Photopigments and circadian systems of vertebrates. Biophys Chem. 1995;

61. Feng J, Yan Z, Ferreira A, Tomizawa K, Liauw JA, Zhuo M, et al. Spinophilin regulates the formation and function of dendritic spines. Proc Natl Acad Sci U S A [Internet]. 2000;97(16):9287–92. Available from: http://www.ncbi.nlm.nih.gov/pubmed/10922077

62. Yang G, Lai CS, Cichon J, Ma L, Li W, Gan WB. Sleep promotes branch-specific formation of dendritic spines after learning. Science (80-) [Internet]. 2014;344(6188):1173–8. Available from: http://www.ncbi.nlm.nih.gov/pubmed/24904169

63. Prince TM, Abel T. The impact of sleep loss on hippocampal function. Learn Mem [Internet]. 2013;20(10):558–69. Available from: http://www.ncbi.nlm.nih.gov/pubmed/24045505

64. Franken P. A role for clock genes in sleep homeostasis. Curr Opin Neurobiol [Internet]. 2013;23(5):864–72. Available from: http://www.ncbi.nlm.nih.gov/pubmed/23756047

65. Xu Y, Padiath QS, Shapiro RE, Jones CR, Wu SC, Saigoh N, et al. Functional consequences of a CKIδ mutation causing familial advanced sleep phase syndrome. Nature. 2005;

66. Franken P, Thomason R, Heller HC, O’Hara BF. A non-circadian role for clock-genes in sleep homeostasis: a strain comparison. BMC Neurosci [Internet]. 2007;8:87. Available from: http://www.ncbi.nlm.nih.gov/pubmed/17945005

67. Moreno-De-Luca A, Helmers SL, Mao H, Burns TG, Melton AM, Schmidt KR, et al. Adaptor protein complex-4 (AP-4) deficiency causes a novel autosomal recessive cerebral palsy syndrome with microcephaly and intellectual disability. J Med Genet [Internet]. 2011;48(2):141–4. Available from: http://www.ncbi.nlm.nih.gov/pubmed/20972249

68. Hofstetter JR, Trofatter JA, Kernek KL, Nurnberger JI, Mayeda AR. New quantitative trait loci for the genetic variance in circadian period of locomotor activity between inbred strains of mice. J Biol Rhythm [Internet]. 2003;18(6):450–62. Available from: http://www.ncbi.nlm.nih.gov/pubmed/14667146

69. Miyake A, Takahashi S, Nakamura Y, Inamura K, Matsumoto S, Mochizuki S, et al. Disruption of the ether-a-go-go K+ channel gene BEC1/KCNH3 enhances cognitive function. J Neurosci [Internet]. 2009;29(46):14637–45. Available from: http://www.ncbi.nlm.nih.gov/pubmed/19923296

70. Zhang X, Bertaso F, Yoo JW, Baumgartel K, Clancy SM, Lee V, et al. Deletion of the potassium channel Kv12.2 causes hippocampal hyperexcitability and epilepsy. Nat Neurosci [Internet]. 2010;13(9):1056–8. Available from: http://www.ncbi.nlm.nih.gov/pubmed/20676103

71. Cirelli C, Bushey D, Hill S, Huber R, Kreber R, Ganetzky B, et al. Reduced sleep in Drosophila Shaker mutants. Nature [Internet]. 2005/04/29. 2005;434(7037):1087–92. Available from: http://www.ncbi.nlm.nih.gov/pubmed/15858564

72. Ding F, O’Donnell J, Xu Q, Kang N, Goldman N, Nedergaard M. Changes in the composition of brain interstitial ions control the sleep-wake cycle. Science (80-) [Internet]. 2016;352(6285):550–5. Available from: http://www.ncbi.nlm.nih.gov/pubmed/27126038

73. Lehotzky A, Lau P, Tokesi N, Muja N, Hudson LD, Ovadi J. Tubulin polymerization-promoting protein (TPPP/p25) is critical for oligodendrocyte differentiation. Glia [Internet]. 2010;58(2):157–68. Available from: http://www.ncbi.nlm.nih.gov/pubmed/19606501

74. Li A, Nattie E. Catecholamine neurones in rats modulate sleep, breathing, central chemoreception and breathing variability. J Physiol [Internet]. 2005/10/29. 2006;570(Pt 2):385–96. Available from: https://www.ncbi.nlm.nih.gov/pubmed/16254009

75. Karp NA, Mason J, Beaudet AL, Benjamini Y, Bower L, Braun RE, et al. Prevalence of sexual dimorphism in mammalian phenotypic traits. Nat Commun [Internet]. 2017/06/27. 2017;8:15475. Available from: https://www.ncbi.nlm.nih.gov/pubmed/28650954

76. Koehl M, Battle S, Meerlo P. Sex differences in sleep: the response to sleep deprivation and restraint stress in mice. Sleep [Internet]. 2006/10/17. 2006;29(9):1224–31. Available from: http://www.ncbi.nlm.nih.gov/pubmed/17040010

77. Paul KN, Dugovic C, Turek FW, Laposky AD. Diurnal sex differences in the sleep-wake cycle of mice are dependent on gonadal function. Sleep [Internet]. 2006/10/17. 2006;29(9):1211–23. Available from: http://www.ncbi.nlm.nih.gov/pubmed/17040009

78. Carrier J, Land S, Buysse DJ, Kupfer DJ, Monk TH. The effects of age and gender on sleep EEG power spectral density in the middle years of life (ages 20-60 years old). Psychophysiology [Internet]. 2001/05/12. 2001;38(2):232–42. Available from: http://www.ncbi.nlm.nih.gov/pubmed/11347869

79. Buysse DJ, Germain A, Hall ML, Moul DE, Nofzinger EA, Begley A, et al. EEG spectral analysis in primary insomnia: NREM period effects and sex differences. Sleep [Internet]. 2008/12/19. 2008;31(12):1673–82. Available from: http://www.ncbi.nlm.nih.gov/pubmed/19090323

80. Collop NA, Adkins D, Phillips BA. Gender differences in sleep and sleep-disordered breathing. Clin Chest Med [Internet]. 2004/04/22. 2004;25(2):257–68. Available from: http://www.ncbi.nlm.nih.gov/pubmed/15099887

81. Krishnan V, Collop NA. Gender differences in sleep disorders. Curr Opin Pulm Med [Internet]. 2006/10/21. 2006;12(6):383–9. Available from: http://www.ncbi.nlm.nih.gov/pubmed/17053485

82. Pavlova M, Sheikh LS. Sleep in women. Semin Neurol [Internet]. 2011/11/25. 2011;31(4):397–403. Available from: http://www.ncbi.nlm.nih.gov/pubmed/22113512

83. Funato H, Miyoshi C, Fujiyama T, Kanda T, Sato M, Wang Z, et al. Forward-genetics analysis of sleep in randomly mutagenized mice. Nature [Internet]. 2016/11/05. 2016;539(7629):378–83. Available from: https://www.ncbi.nlm.nih.gov/pubmed/27806374

84. Hayasaka N, Hirano A, Miyoshi Y, Tokuda IT, Yoshitane H, Matsuda J, et al. Salt-inducible kinase 3 regulates the mammalian circadian clock by destabilizing PER2 protein. Elife [Internet]. 2017/12/12. 2017;6. Available from: https://www.ncbi.nlm.nih.gov/pubmed/29227248

