## Supplementary Data for "Noninvasive sleep monitoring in large-scale screening of knock-out mice reveals novel sleep-related genes"

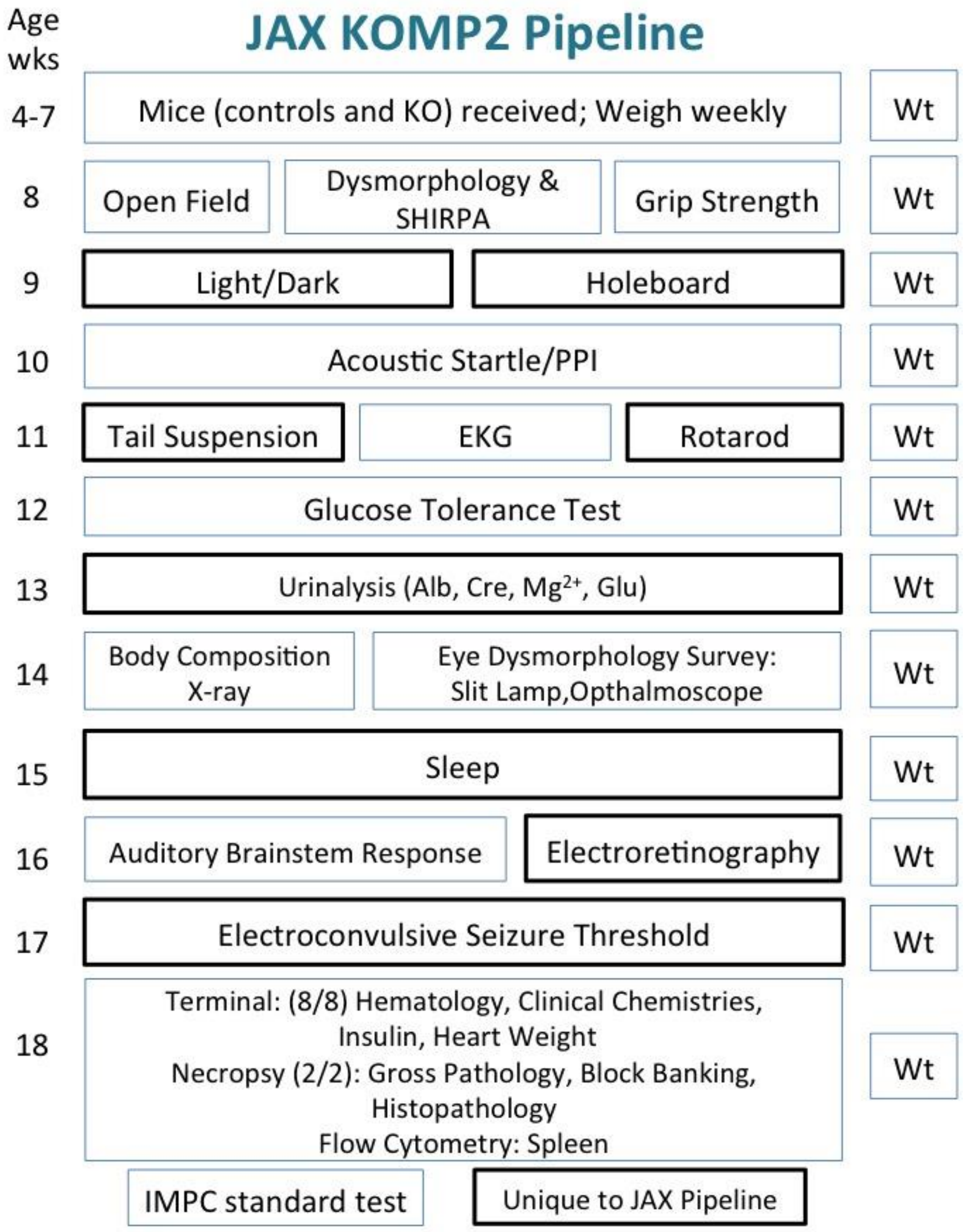


Supplementary Fig 1: Phenotyping pipeline at The Jackson Laboratory used to evaluate mice included in this report.

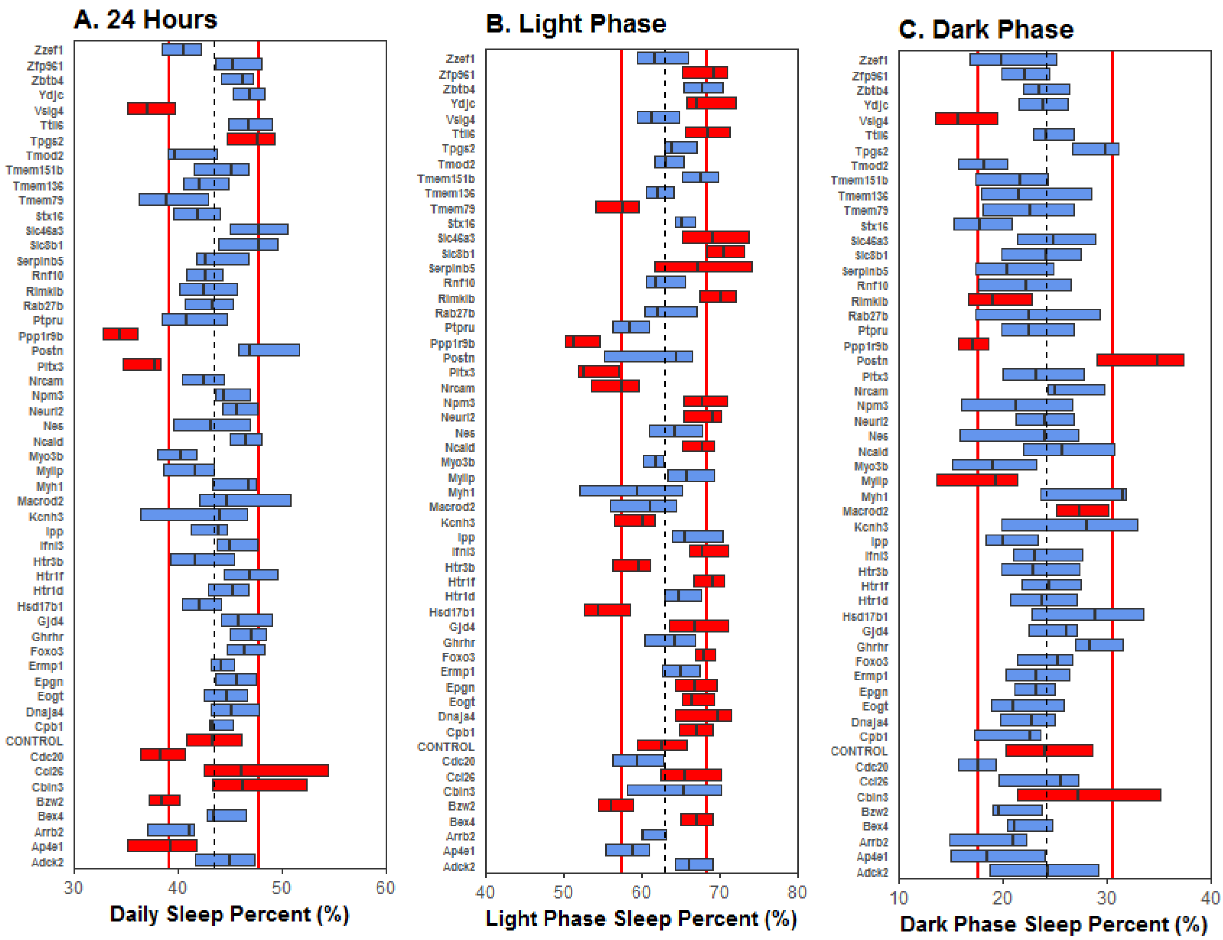
A. 24 Hours
Zzef1
Zfp961
Zbtb4
Ydjc
Vsig4
Ttll6
Tpgs2
Tmod2
Tmem151b
Tmem136
Tmem79
Stx16
Slc46a3
Slc8b1
Serpinb5
Rnf10
Rimklb
Rab27b
Ptpru
Ppp1r9b
Postn
Pitx3
Nrcam
Npm3
Neurl2
Nes
Ncald
Myo3b
Mylip
Myh1
Macrod2
Kcnh3
Ipp
Ifnl3
Htr3b
Htr1f
Htr1d
Hsd17b1
Gjd4
Ghrhr
Foxo3
Ermp1
Epgn
Eogt
Dnaja4
Cpb1
CONTROL
Cdc20
Ccl26
Cbln3
Bzw2
Bex4
Arrb2
Ap4e1
Adck2
30
40
50
60
Daily Sleep Percent (%)
B. Light Phase
40
50
60
70
80
Light Phase Sleep Percent (%)
C. Dark Phase
10
20
30
40
Dark Phase Sleep Percent (%)

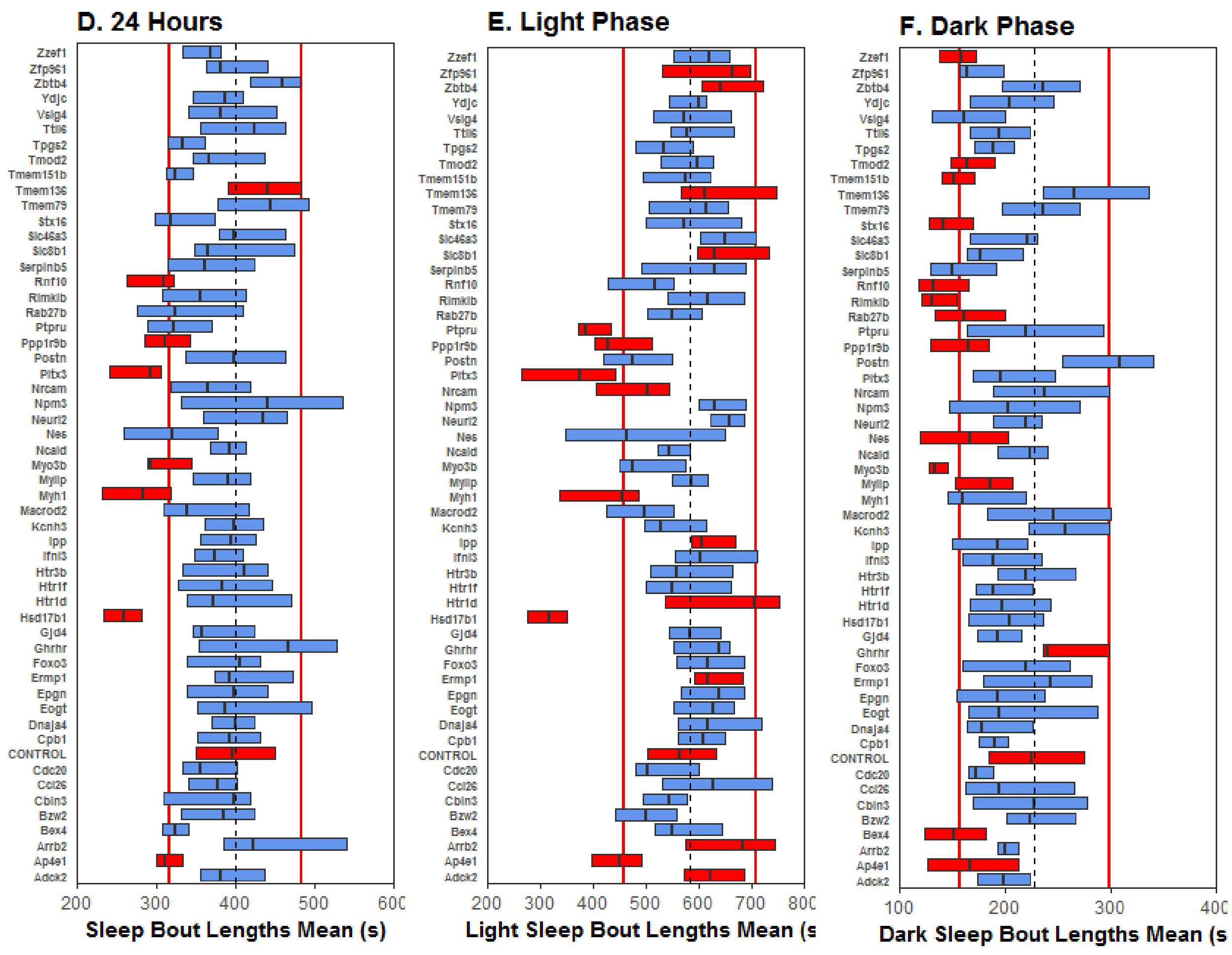


Supplementary Figure 2: Visual representation of the differences in values for sleep variables in 55 KO strains that were found to be significant for one or more of these variables. The genotypes are arranged alphabetically for sleep percent (A. 24h, B. light phase, C. dark phase), and bout length (D. 24h, E. light phase, F. dark phase) measured in seconds as depicted on the x axis. Each bar represents values between the first and third quantile and the median (dotted line) for each KO cohort. Red boxes represent the genes significant for a specific PiezoSleep variable.

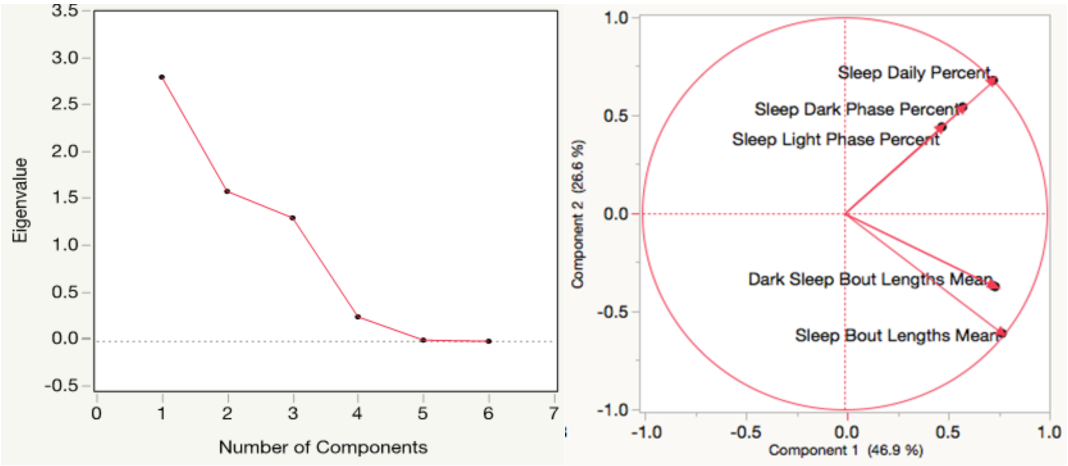


A. Scree Plot B. Biplot

Supplementary Figure 3: Scree plot depicting relationship between eigenvalue and number of components.

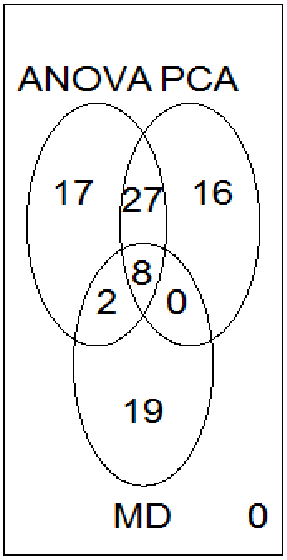


Figure 4: Venn diagram representing number of genes from ANOVA, PCA and Mahalanobis distance outliers.

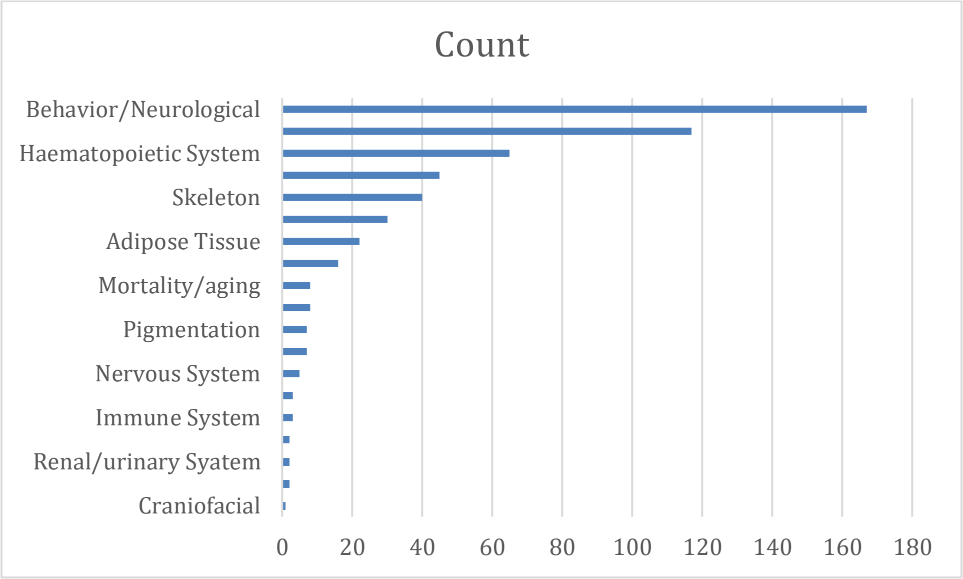


Figure 5: Frequency of Mammalian Phenotype (MP) ontology in the 122 sleep related candidate genes.

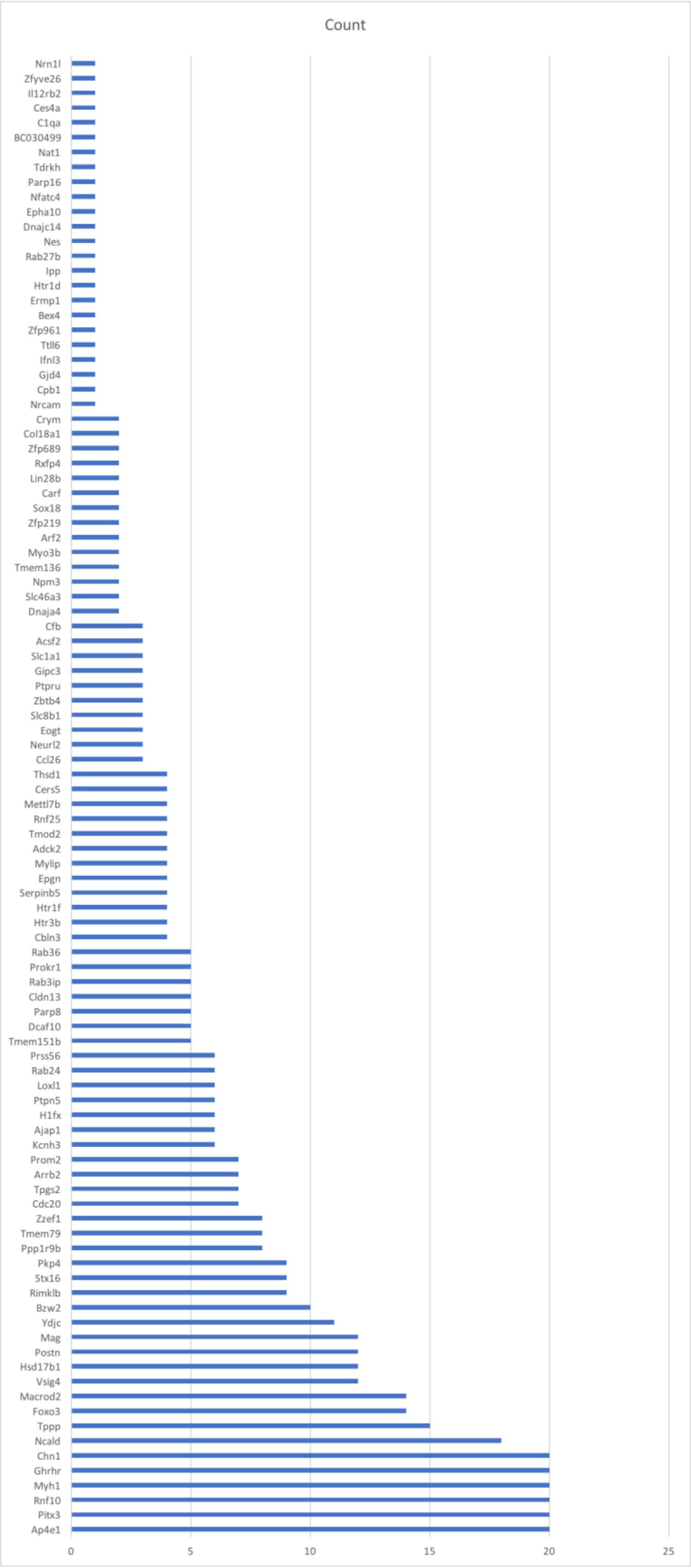
Count
Nrn1l
Zfyve26
Il12rb2
Ces4a
C1qa
BC030499
Nat1
Tdrkh
Parp16
Nfatc4
Epha10
Dnajc14
Nes
Rab27b
Ipp
Htr1d
Ermp1
Bex4
Zfp961
Ttll6
Ifnl3
Gjd4
Cpb1
Nrcam
Crym
Col18a1
Zfp689
Rxfp4
Lin28b
Carf
Sox18
Zfp219
Arf2
Myo3b
Tmem136
Npm3
Slc46a3
Dnaja4
Cfb
Acsf2
Slc1a1
Gipc3
Ptpru
Zbtb4
Slc8b1
Eogt
Neurl2
Ccl26
Thsd1
Cers5
Mettl7b
Rnf25
Tmod2
Adck2
Mylip
Epgn
Serpinb5
Htr1f
Htr3b
Cbln3
Rab36
Prokr1
Rab3ip
Cldn13
Parp8
Dcaf10
Tmem151b
Prss56
Rab24
Loxl1
Ptpn5
H1fx
Ajap1
Kcnh3
Prom2
Arrb2
Tpgs2
Cdc20
Zzef1
Tmem79
Ppp1r9b
Pkp4
Stx16
Rimklb
Bzw2
Ydjc
Mag
Postn
Hsd17b1
Vsig4
Macrod2
Foxo3
Tppp
Ncald
Chn1
Ghrhr
Myh1
Rnf10
Pitx3
Ap4e1
0
5
10
15
20
25

Figure 6: Mammalian Phenotype (MP) terms were associated with 122 genes. Each gene shown had from 1-20 associated MP terms. Genes not listed (n=22) were not associated with MP terms.

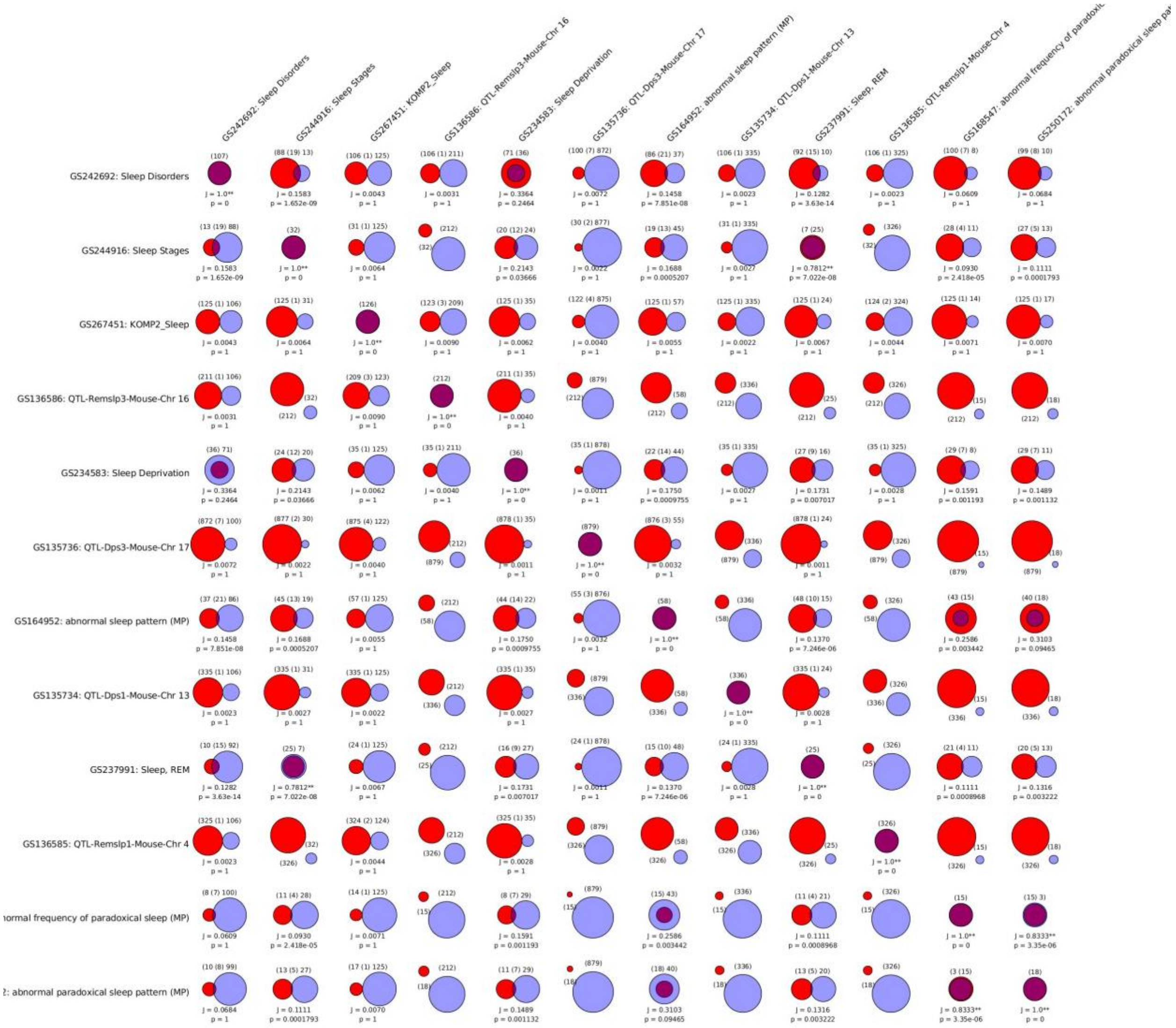


Figure 7: Jaccard Analysis for geneset similarities from 12 shortlisted genelists from Geneweaver.

**Supplementary Tables:**

### Loading Matrix

| | Prin1 | Prin2 | Prin3 | Prin4 | Prin5 | Prin6 |
|---|---|---|---|---|---|---|
| Sleep Bout Lengths Mean | 0.77928 | -0.61304 | 0.04154 | 0.09501 | -0.07840 | 0.00000 |
| Light Sleep Bout Lengths Mean | 0.73327 | -0.36842 | 0.45257 | -0.34714 | 0.03541 | -0.00000 |
| Dark Sleep Bout Lengths Mean | 0.74518 | -0.37461 | -0.50148 | 0.22108 | 0.06341 | -0.00000 |
| Sleep Daily Percent | 0.73559 | 0.67732 | -0.00028 | 0.00840 | -0.00815 | -0.00042 |
| Sleep Light Phase Percent | 0.48215 | 0.44026 | 0.72222 | 0.22800 | 0.01080 | 0.00026 |
| Sleep Dark Phase Percent | 0.58471 | 0.54134 | -0.57900 | -0.17161 | -0.01939 | 0.00032 |

**Table 1:** ***Loading matrix for PCs***

| Genotype | Per1 | Per2 | Clock | Bmal1/Arntl | Homer1 | Sik3 | Nalcn |
|---|---|---|---|---|---|---|---|
| Ap4e1 | 1 | 0 | 0 | 0 | 0 | 0 | 0 |
| BC030499 | 0 | 0 | 0 | 0 | 0 | 1 | 0 |
| Bex4 | 1 | 0 | 0 | 0 | 0 | 0 | 0 |
| Cacna2d3 | 1 | 0 | 1 | 1 | 0 | 0 | 1 |
| Chn1 | 0 | 0 | 0 | 0 | 1 | 0 | 0 |
| Dcaf10 | 0 | 0 | 1 | 0 | 0 | 0 | 0 |
| Dnaja4 | 0 | 0 | 1 | 0 | 0 | 0 | 0 |
| Eogt | 1 | 0 | 0 | 0 | 0 | 0 | 0 |
| Epgn | 0 | 0 | 0 | 1 | 0 | 0 | 0 |
| Ermp1 | 0 | 1 | 0 | 0 | 0 | 0 | 0 |
| Foxo3 | 0 | 1 | 0 | 0 | 0 | 0 | 0 |
| H1fx | 1 | 0 | 0 | 0 | 0 | 0 | 0 |
| Htr1f | 0 | 0 | 0 | 1 | 0 | 0 | 0 |
| Igsf11 | 1 | 0 | 0 | 0 | 0 | 0 | 0 |
| Ipp | 0 | 0 | 0 | 1 | 0 | 0 | 0 |
| Kcnh3 | 1 | 1 | 1 | 1 | 0 | 0 | 0 |
| Loxl1 | 1 | 0 | 0 | 0 | 0 | 0 | 0 |

| | | | | | | | |
|---|---|---|---|---|---|---|---|
| Mos | 0 | 0 | 0 | 0 | 0 | 1 | 0 |
| Myo3a | 0 | 0 | 0 | 1 | 0 | 0 | 1 |
| Myo3b | 0 | 0 | 0 | 0 | 1 | 0 | 0 |
| Nat1 | 0 | 0 | 0 | 0 | 1 | 0 | 0 |
| Ncald | 0 | 0 | 0 | 0 | 0 | 0 | 1 |
| Nek2 | 0 | 0 | 0 | 0 | 0 | 1 | 0 |
| Nes | 1 | 0 | 0 | 0 | 0 | 0 | 0 |
| Nrcam | 0 | 0 | 1 | 0 | 0 | 0 | 0 |
| Parp16 | 0 | 0 | 0 | 0 | 1 | 0 | 0 |
| Parp8 | 0 | 0 | 0 | 1 | 0 | 0 | 0 |
| Pkp4 | 0 | 1 | 0 | 0 | 0 | 0 | 0 |
| Ppp1r9b | 1 | 0 | 0 | 0 | 0 | 0 | 0 |
| Prom1 | 0 | 1 | 1 | 0 | 0 | 0 | 0 |
| Ptpn5 | 0 | 0 | 0 | 0 | 0 | 0 | 1 |
| Ptpru | 1 | 0 | 0 | 0 | 0 | 0 | 0 |
| Rab27b | 0 | 0 | 1 | 0 | 0 | 0 | 0 |
| Rab36 | 0 | 0 | 0 | 0 | 0 | 0 | 1 |
| Rnf25 | 1 | 0 | 0 | 0 | 0 | 0 | 0 |
| Slc46a3 | 0 | 1 | 0 | 0 | 0 | 0 | 0 |
| Srcin1 | 0 | 0 | 0 | 0 | 0 | 0 | 1 |
| Stx16 | 0 | 0 | 0 | 0 | 0 | 1 | 0 |
| Tpgs2 | 0 | 0 | 1 | 1 | 0 | 0 | 0 |
| Tppp | 0 | 0 | 0 | 0 | 1 | 0 | 1 |

Table 2: Binary table representing connections in gene network analysis. 1 indicates that gene is part of the network.

| MP Term | Count |
|---|---|
| abnormal sleep behavior | 37 |
| abnormal behavioral response to light | 23 |
| decreased circulating glucose level | 22 |
| Info not available | 22 |
| decreased total body fat amount | 20 |
| abnormal behavior | 16 |
| hyperactivity | 14 |
| abnormal bone structure | 14 |
| decreased bone mineral content | 14 |
| decreased lean body mass | 12 |
| increased circulating sodium level | 11 |
| decreased bone mineral density | 11 |
| increased circulating alkaline phosphatase level | 10 |
| decreased circulating insulin level | 10 |
| decreased grip strength | 10 |
| increased lean body mass | 9 |
| decreased body length | 8 |
| increased hematocrit | 8 |
| increased circulating potassium level | 8 |
| impaired righting response | 8 |
| increased fasted circulating glucose level | 7 |

| | |
|---|---|
| hypoactivity | 7 |
| convulsive seizures | 7 |
| abnormal retinal pigmentation | 7 |
| increased bone mineral content | 6 |
| decreased leukocyte cell number | 6 |
| increased mean corpuscular volume | 6 |
| increased mean corpuscular hemoglobin | 6 |
| increased hemoglobin content | 6 |
| decreased circulating triglyceride level | 6 |
| abnormal retina morphology | 6 |
| abnormal QRS complex | 5 |
| increased erythrocyte cell number | 5 |
| improved glucose tolerance | 5 |
| decreased circulating free fatty acid level | 5 |
| abnormal coat appearance | 5 |
| shortened RR interval | 4 |
| increased body length | 4 |
| increased bone mineral density | 4 |
| increased or absent threshold for auditory brainstem response | 4 |
| preweaning lethality, incomplete penetrance | 4 |
| increased exploration in new environment | 4 |
| decreased vertical activity | 4 |
| increased grip strength | 4 |
| decreased exploration in new environment | 4 |

| | |
|---|---|
| increased vertical activity | 4 |
| decreased startle reflex | 4 |
| decreased anxiety-related response | 4 |
| persistence of hyaloid vascular system | 4 |
| decreased heart weight | 3 |
| cardiovascular system phenotype | 3 |
| decreased mean corpuscular hemoglobin concentration | 3 |
| thrombocytopenia | 3 |
| decreased threshold for auditory brainstem response | 3 |
| increased circulating HDL cholesterol level | 3 |
| decreased circulating HDL cholesterol level | 3 |
| decreased circulating cholesterol level | 3 |
| decreased circulating chloride level | 3 |
| decreased prepulse inhibition | 3 |
| increased total body fat amount | 2 |
| increased heart rate | 2 |
| shortened PQ interval | 2 |
| increased heart weight | 2 |
| increased mean corpuscular hemoglobin concentration | 2 |
| decreased mean corpuscular hemoglobin | 2 |
| increased blood urea nitrogen level | 2 |
| increased circulating glucose level | 2 |
| decreased fasted circulating glucose level | 2 |
| increased circulating chloride level | 2 |

| | |
|---|---|
| decreased blood urea nitrogen level | 2 |
| enlarged lymph nodes | 2 |
| abnormal coat/hair pigmentation | 2 |
| increased prepulse inhibition | 2 |
| increased coping response | 2 |
| straub tail | 2 |
| limb grasping | 2 |
| abnormal motor coordination/ balance | 2 |
| abnormal kidney morphology | 2 |
| male infertility | 2 |
| decreased heart rate variability | 1 |
| abnormal heart morphology | 1 |
| shortened ST segment | 1 |
| abnormal retinal blood vessel morphology | 1 |
| prolonged QRS complex duration | 1 |
| thin ventricular wall | 1 |
| shortened PR interval | 1 |
| prolonged PQ interval | 1 |
| prolonged ST segment | 1 |
| absent teeth | 1 |
| increased body weight | 1 |
| abnormal head morphology | 1 |
| enlarged spleen | 1 |
| increased gamma-delta T cell number | 1 |

| | |
|---|---|
| decreased alpha-beta T cell number | 1 |
| increased CD4-positive, CD25-positive, alpha-beta regulatory T cell number | 1 |
| decreased erythrocyte cell number | 1 |
| increased memory CD4-positive, CD25-positive, alpha-beta regulatory T cell number | 1 |
| decreased Ly6C-positive NK T cell number | 1 |
| increased marginal zone B cell number | 1 |
| decreased hemoglobin content | 1 |
| increased leukocyte cell number | 1 |
| decreased mean corpuscular volume | 1 |
| decreased memory-marker CD4-negative NK T cell number | 1 |
| decreased effector memory T-helper cell number | 1 |
| increased CD8-positive, naive alpha-beta T cell number | 1 |
| decreased macrophage cell number | 1 |
| increased CD8-positive, alpha-beta T cell number | 1 |
| decreased basophil cell number | 1 |
| decreased eosinophil cell number | 1 |
| decreased circulating amylase level | 1 |
| increased circulating alanine transaminase level | 1 |
| increased circulating bilirubin level | 1 |
| decreased circulating serum albumin level | 1 |
| decreased circulating alkaline phosphatase level | 1 |
| decreased urine creatinine level | 1 |
| increased circulating phosphate level | 1 |
| increased circulating iron level | 1 |

| | |
|---|---|
| decreased circulating iron level | 1 |
| decreased circulating phosphate level | 1 |
| decreased circulating alanine transaminase level | 1 |
| immune system phenotype | 1 |
| abnormal nail morphology | 1 |
| short tail | 1 |
| abnormal digit morphology | 1 |
| preweaning lethality, complete penetrance | 1 |
| embryonic lethality prior to tooth bud stage | 1 |
| embryonic lethality prior to organogenesis | 1 |
| prenatal lethality prior to heart atrial septation | 1 |
| abnormal social/conspecific interaction | 1 |
| impaired pupillary reflex | 1 |
| jumpy | 1 |
| absent startle reflex | 1 |
| abnormal vocalization | 1 |
| abnormal motor learning | 1 |
| decreased coping response | 1 |
| increased startle reflex | 1 |
| tremors | 1 |
| abnormal whole-body plethysmography | 1 |
| increased tidal volume | 1 |
| decreased pulmonary respiratory rate | 1 |
| increased sacral vertebrae number | 1 |

| | |
|---|---|
| anophthalmia | 1 |
| abnormal eyelid aperture | 1 |
| mydriasis | 1 |
| abnormal lens morphology | 1 |
| decreased cornea thickness | 1 |
| fused cornea and lens | 1 |

Supplementary Table 3: Column “Count” represents the number of times a Mammalian Phenotype (MP) term was associated with a gene.
